# Retinal adaptive mechanisms confer selectivity to homogeneous objects in natural scenes

**DOI:** 10.64898/2026.09.22.753530

**Authors:** Baptiste Lorenzi, Déborah Varro, Rémi Baroux, Samuele Virgili, Olivier Marre

## Abstract

Adaptive mechanisms in sensory neurons are crucial to transmit information in different contexts. In the retina, it is assumed that their role is to normalize neuronal responses to input statistics like mean and variance. However, this role has mostly been characterized with simple, artificial stimuli, and remains unclear for natural stimuli. Here we show that during their response to natural scenes, adaptive mechanisms reshape the feature selectivity of ganglion cells, the retinal output. We recorded retinal ganglion cell responses to rapid sequences of natural images in mice. Including a bio-inspired adaptive mechanism in an artificial neural network model was necessary to predict cell responses to new sequences of natural images. This adaptive mechanism tuned specific cell types to selectively respond to homogeneous regions situated within cluttered visual surrounds, a feature suited for detecting threats. Adaptive mechanisms do not merely normalize responses but actively enable new feature selectivity in the early visual system.

## Introduction

Our visual system allows us to achieve tasks robustly in many different visual contexts, coping with an ever changing sensory input, despite having a limited bandwidth to do so. For example, we can experience changes of luminance across 14 orders of magnitude, while our Retinal ganglion cells (RGCs) can modulate their firing rate only across 2 or 3 orders of magnitude. A primary solution to this challenge is physiological adaptation: a dynamic modulation of the neural code based on recent sensory history^1^. Adaptation can happen over different time scales: slow adaptation is a gradual process that can take several seconds^2,3^. Short term adaptation occurs in less than a second^3,4^.

Several mechanisms are thought to form the biological substrate of this short-term adaptation. Synaptic weight decreases following a previous vesicle release, making synaptic weight history dependent. Neuronal intrinsic properties, like inactivating ion channels, can confer adaptive properties to single neurons. Inhibitory circuits, e.g. feedforward inhibition, can also make neuronal responses dependent on the previous stimulus history.

These adaptive mechanisms will change how a neuron responds to a visual stimulus depending on the visual context, and will allow us to perform visual tasks in different visual contexts. A model that aims at replicating how a sensory circuit responds to visual stimuli in different contexts should thus account for these mechanisms. Understanding how we can process stimuli in various contexts requires modeling these adaptive mechanisms, and understanding their functional impact.

Classically, in the retina, it is assumed that their functional role is to normalize the responses with respect to the input statistics, subtracting the mean and dividing by the standard deviation^4^. Adaptation to the mean intensity of a scene has been studied extensively and already happens at the level of photoreceptors^5–7^. The retina can also adapt to the standard deviation of light intensity (i.e. contrast), either over space or time, by changing the gain of the response^5,8^. As a consequence, the retina can be viewed as a feature extractor whose dynamic range is maintained by adaptive normalization mechanisms. Artificial neural networks typically reproduce this organization, where each layer is composed of a feature extraction stage followed by a normalization.

However, adaptive mechanisms have been characterized and modeled as a normalization step mostly in stimulus conditions that are pretty far from natural inputs. The complex natural statistics may engage adaptation mechanisms in fundamentally different ways than for simple gratings or checkerboards^6^. Other forms of adaptation like sensitization, a transient increase of gain following an increase in stimulus standard deviation^9,10^; adaptation to higher order statistics^11^ or pattern adaptation^12^, suggest that adaptive mechanisms may play a more complex role, but they often have been studied with stimuli that are pretty far from natural scenes. In response to natural stimuli, models composed of feature maps and normalization steps, like deep networks, can have a hard time to generalize and predict retinal response to new stimulus statistics. Including detailed biophysical models of adaptation processes can help to overcome this generalization problem^13^, suggesting that a pure normalization may not entirely reproduce their functional impact. This raises the question of whether adaptive mechanisms can still be modelled as a normalization when neurons respond to a natural scene instead of artificial stimuli.

Here we show that during natural scene stimulation, adaptive mechanisms do not just act to normalize the response, but also reshape the feature selectivity of retinal ganglion cells. We characterize the selectivity of mouse ganglion cells during natural scene stimulation, and how it is reshaped by the recent stimulus history. We found that spatial, as well as ON/OFF selectivity, can change with stimulus history. We designed a model that could reproduce this history dependence. A key mechanism that allowed this model to generalize and predict responses to unseen stimulus history was a biologically plausible gain control. The model predicted that a specific type of ganglion cells, the W3 cells, should be selective for large, uniform objects embedded in a cluttered natural background. Gain control was essential for this selectivity. We confirmed experimentally that this was the case, demonstrating that specific types of cells have a function that can be directly useful for the detection of threats embedded in natural scenes. This shows that adaptive mechanisms are not just useful to adapt neural responses to the global statistics of the visual input, but also reshape the selectivity of sensory neurons.

## Results

### Ganglion cells can change their selectivity depending on short term stimulus history

We recorded ganglion cells in the mouse retina using multi-electrode arrays while stimulating photoreceptors with a combination of artificial and natural images. To probe short term adaptation in retinal ganglion cells, we displayed paired image sequences selected from a large database of natural images^14^: an adaptor for 400 ms and a probe for 400 ms, together forming an adaptor-probe sequence lasting 800ms (Fig. 1A). Notably, the duration of each image here is similar to the average interval between two saccades in the mouse^15^. We found that the adaptor modulated the response of cells to the probe (Fig. 1B). To assess the total magnitude of this modulation, we defined an adaptation index for each neuron as the relative change in firing rate compared to the non-adapted baseline (see Methods). Across the population, this adaptation index was on average 67% ± 38% (mean ± SD, n = 325 cells; Wilcoxon signed-rank test against 0, p < 0.001) confirming that a single probe can generate strikingly different responses for two different adaptors.

**Fig. 1.**
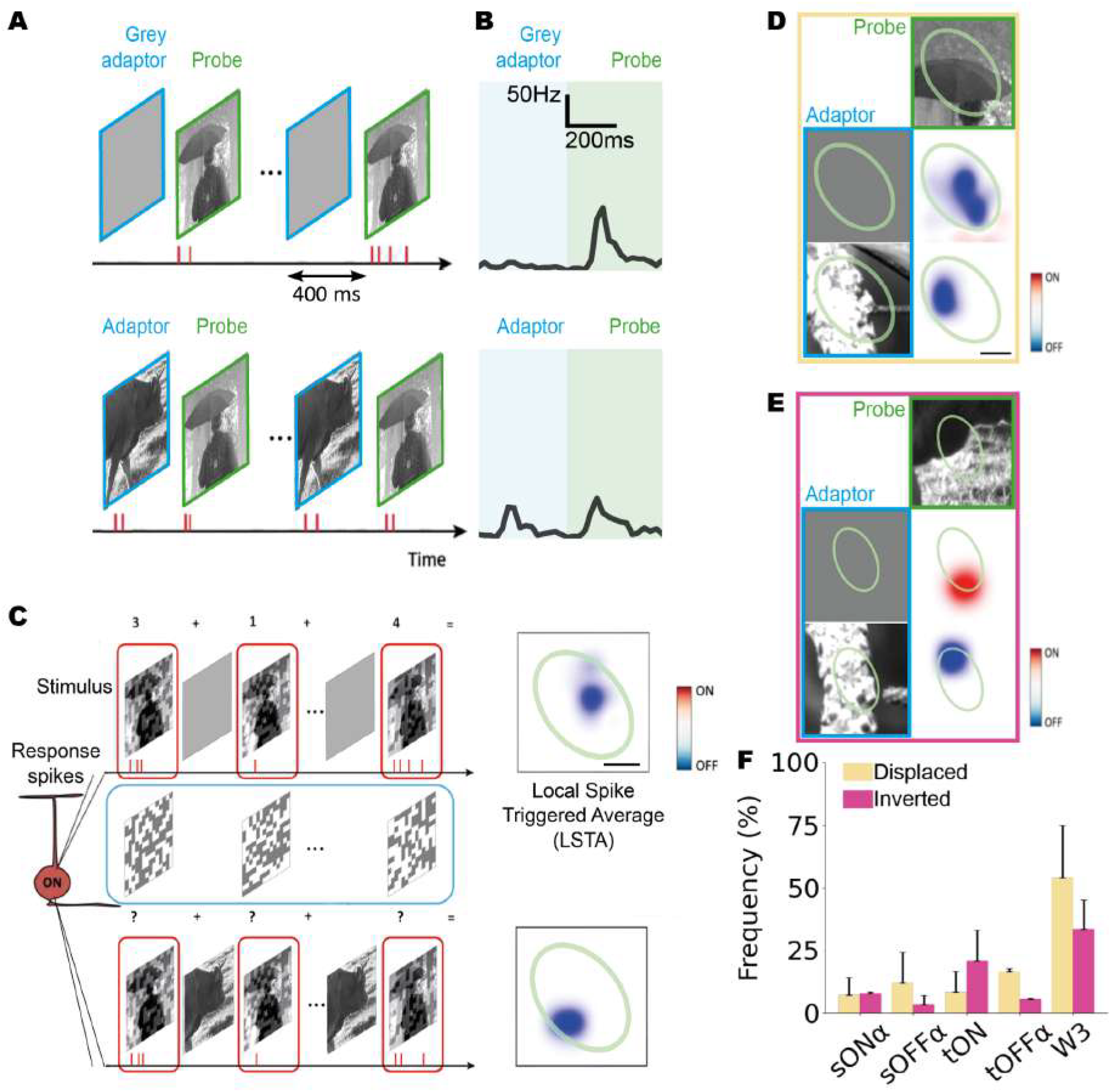
Ganglion cells change their selectivity to natural scenes depending on short-term stimulus history. **A**. Schematic of the visual stimuli : two images were presented successively for each sequence. Each image was presented for 400ms. In each sequence, a first natural image (probe) was preceded by another image (adaptor) : grey (to prevent adaptation) or another natural image. **B**. Peri-Stimulus Time Histograms (PSTHs) showing the response of one example cell to the two stimuli shown in A. Responses are averaged over 30 repetitions (see Methods). **C**. We presented a randomized combination of the adaptor-probe sequences described in A, but using perturbed probes (second image in the sequence) and compared how LSTA changed depending on the probe. To calculate the LSTAs, for every probe we calculated a weighted average of the different perturbations by the number of spikes they evoked, independently for each adaptor-probe sequence. Scale bar: 100 µm. **D**. Two example LSTAs, measured for an example cell and perturbed probe paired with two different adaptors. A yellow ellipse fitted to the spatial component of the classical receptive field is shown in all the bottom panels as a reference. Recorded LSTAs were denoised using a Gaussian Process denoiser before plotting^21^ (see Methods). The LSTA is displaced by the presence of the adaptor in the sequence. Left: adaptation image. Top row : Probes, Left column: Adaptors, Bottom right : LSTA measured for the adaptor-probe sequence. Scale bar: 100 µm. **E**. Same as D for another cell and probe. The polarity of the LSTA is inverted due to the presence of the adaptor in the sequence. **F**. Percentage of retinal ganglion cell types showing displaced or inverted LSTAs for 5 different cell types (n = 68 sustained-ONα cells (sONα), 32 sustained-OFFα cells (sOFFα), 27 transient-ON cells (tON), 36 transient-OFFα cells (tOFFα), 29 W3 cells). Data are presented as mean and SEM.

While the changes in ganglion cells’ responses confirm they are affected by short term adaptation, it remains unclear if this adaptation reshapes their selectivity as well. To address this issue we estimated the selectivity of ganglion cells during natural stimulation using the perturbative approach described in Goldin et al.^16^. We added a low-contrast checkerboard superposed on the probes. Each different checkerboard created a small but measurable change in the cells’ responses compared to the unperturbed image case. To determine which perturbation would evoke the largest increase in response, for every cell and for every adaptor-probe sequence we averaged the perturbations weighted by the number of elicited spikes (Fig. 1C). This is similar to the classical Spike-Triggered Average (STA) analysis but exploring sensitivity to small stimulus perturbations around the probe. We therefore refer to them as Local Spike-Triggered Averages (LSTAs). For instance, an ON-type LSTA indicates that a luminance increase is the best way to elicit more spikes in the context of a given image.

We observed that two LSTAs measured for the same probe could be vary for different adaptors (see Methods), confirming short term adaptation can reshape ganglion cell’s selectivity. This reshaping most often appeared as a displacement of the LSTA: for a same probe, different adaptors led to different spatial selectivity (Fig. 1D). We also observed changes in polarity selectivity (from ON to OFF or viceversa) due to a change in adaptor (Fig. 1E). These results demonstrate that the ON-OFF selectivity of retinal ganglion cells in the context of one natural image is not fixed but it is a function of short-term stimulus history. To evaluate the prevalence of this context dependence across retinal ganglion cells, we classified the recorded ganglion cells using standard methods, based on drifting gratings and chirp stimuli^17^ (see Methods). By analyzing the displacement of the center of LSTA, we found that W3 cells were much more likely than other types to exhibit spatial shifts due to adaptation (Fig. 1F, W3: 54.2 ± 29.5%, n = 2 experiments, see Methods). Similarly, when characterizing polarity inverting cells based on LSTA sign changes, W3 cells showed a higher prevalence of inversion compared to other cell types (Fig. 1F, W3: 33.6 ± 16.1%, n = 2 experiments, see Methods).

Overall, these results show that fast adaptive mechanisms do not provide a global normalization of retinal ganglion cells responses alone but also change their spatial and luminance selectivity depending on the recent stimulus history.

### A convolutional neural network with a gain control mechanism predicts the effect of the adaptor on neural response

To explain the modulation of responses and selectivities of retinal ganglion cells due to adaptation, we looked for a model that captured the underlying adaptation mechanism. Such a model should be able to generalize to unseen adaptor-probe sequences, predicting how the response is modulated by adaptors it has not encountered during training. To this end, we trained models exclusively on pairs with a grey adaptor, ensuring no adaptation occurred during training. We then tested whether these models could generalize to new sequences by presenting various adaptors in the test set.

Two-layer convolutional neural networks (CNNs) have been shown to possess the necessary expressivity to model retinal responses to natural stimuli^16,18–21^. We therefore implemented a “Core” CNN to capture neural responses to flashed natural images (Fig. 2A). In this architecture, two convolutional filters (one ON, one OFF at training initialization) extract local spatio-temporal features, which are then integrated by a readout layer to predict firing rates (see Methods). We first evaluated the Core CNN’s ability to predict responses to unseen natural images shown after a grey adaptor. This control set (Fig. 1A) shares the same statistical distribution as the training data (In-Distribution). Qualitatively, the model successfully reproduced the PSTHs recorded across the population (Fig. 2B, see Methods). We then investigated whether these models could generalize to unseen temporal contexts, where the adaptor was a natural image rather than a grey screen (Fig. 2A, second stimulus). In this out-of-distribution (OOD) regime, the Core CNN performed poorly, exhibiting a tendency to overshoot the firing rate immediately following the adaptor (Fig. 2B, bottom, Fig. 2C).

**Fig. 2.**
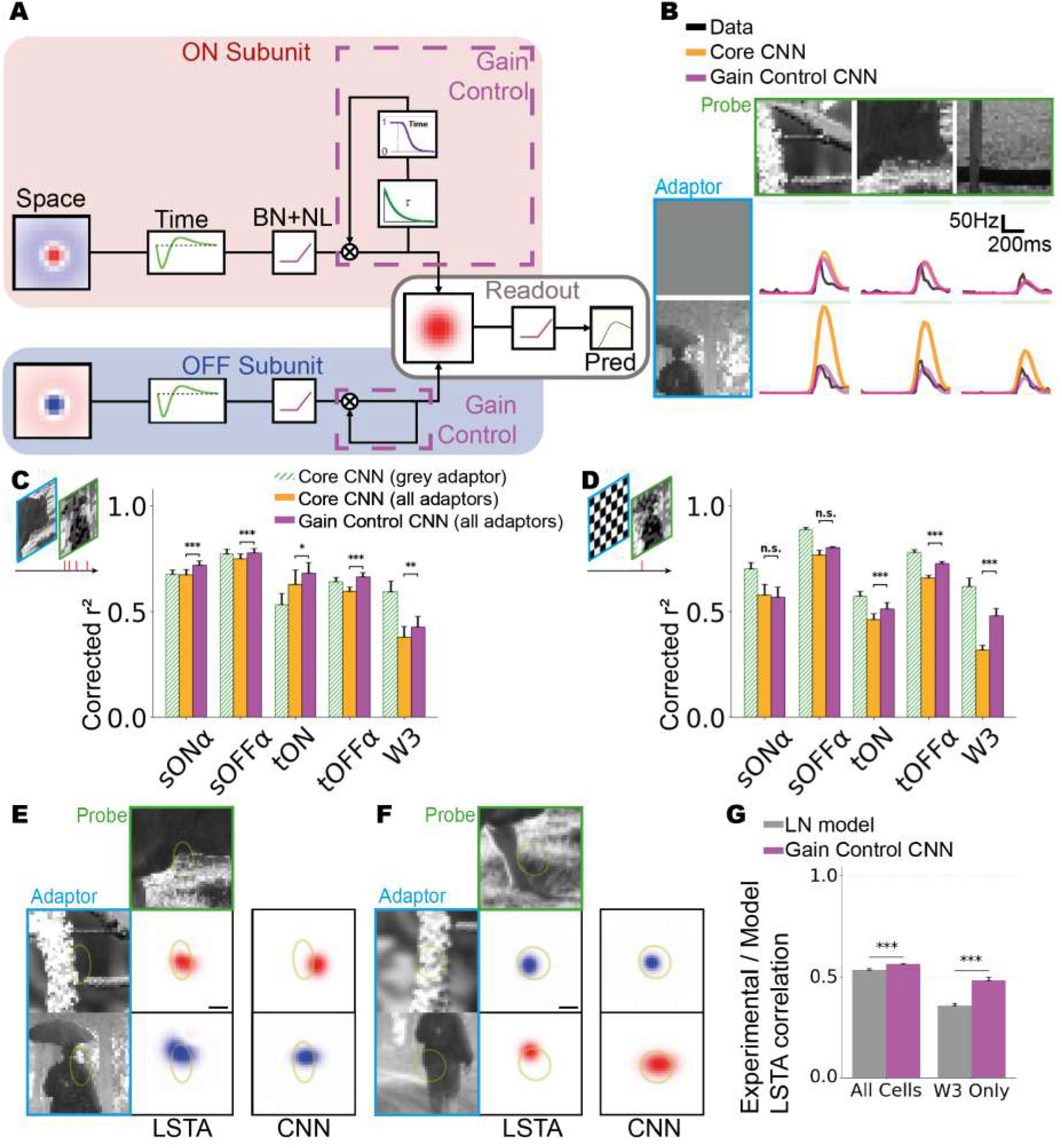
A convolutional neural network with a gain control mechanism predicts the effect of the adaptor on neural response. **A**. Schematic of the different architectures used to predict the response of a single retinal ganglion cell. The core model is composed of an ON Subunit, an OFF Subunit and a Spatial Readout blocks. The subunit blocks are composed of a spatio-temporal convolutional filter, Batch Normalization and a non linearity (BN+NL). The readout is composed of a spatial convolutional filter, Batch Normalization, feature weights and a non linearity. The Gain Control CNN is constructed by adding to the core model a gain control layer in the ON and OFF subunits (Methods). **B**. Example recorded and predicted PSTHs by the two models for one example cell. Top row: Probe, Left column: Adaptor, Bottom right : recorded and predicted PSTHs **C**. Performance of the two models at predicting responses to repeated adaptor-probe sequences with unperturbed probes (see Methods). Only cells that met our reliability criterion and on which we were able to train the Gain Control CNN are presented here (mean reliability > 0.3, see Methods). Bars represent the population mean, and error bars indicate the standard error of the mean (SEM). The Gain Control CNN model significantly outperformed the Core CNN across all cell types (two-sided paired Wilcoxon signed-rank test): sustained-ONα (n = 60 cells, p < 1×10^−4^), sustained-OFFα (n = 25 cells, p < 1×10^−4^), transient-ON (n = 12 cells, p = 0.031), transient-OFFα (n = 31 cells, p < 1×10^−4^), and W3 (n = 15 cells, p = 0.0015). **D**. Same as C but for another set of cells for which the adaptors were checkerboards. The Gain Control CNN model significantly outperformed the Core CNN for some cell types (two-sided paired Wilcoxon signed-rank test): transient-ON (n = 38 cells, p = 0.0001), transient-OFFα (n = 38 cells, p < 1×10^−4^), and W3 (n = 23 cells, p < 1×10^−4^) but not for sustained-ONα (n = 4 cells, p = 0.38), sustained-OFFα (n = 3 cells, p = 0.5). **E**. Comparison of an experimental LSTA with the prediction from the Gain Control CNN for an example cell. Scale bar: 100 µm. **F**. Same as E for another example cell. **G**. Average performance of the LN model and the Gain Control CNN model at predicting the measured LSTAs (see Methods). The Gain Control CNN significantly outperforms the LN (two-sided paired Student’s t-test), in all cell types (n = 1746 LSTAs, p < 1×10^−4^) and in W3 cells (n = 180 LSTAs, p < 1×10 4). Data are presented as mean and SEM.

We then asked whether incorporating bio-inspired adaptive mechanisms in the CNN model could improve the capacity of the model to generalize to such new adaptors. To test this we developed a Gain Control CNN by incorporating a gain control layer following the subunit non-linearity (independently for each subunit). Inspired by Chen et al.^22^, this layer utilizes divisive normalization to dynamically rescale the subunit’s output feature maps based on recent subunit activity (Fig. 2A; see Methods). The Gain control CNN significantly improved the prediction accuracy in the OOD regime for all cell types (Fig. 2B, C).

Finally, we used a high-contrast checkerboard adaptor followed by natural probes to further drive gain modulation (see Methods). In this high-contrast regime, explicit gain control was essential for accurate prediction. While the Core CNN’s performance collapsed on samples including checkerboard adaptors followed by natural probes, the Gain Control CNN maintained high predictive power (Fig. 2D). Specifically, W3 cells, which were largely missed by the Core CNN, were well-captured by the Gain Control CNN (Core: R^2^ = 0.176 ± 0.12; Gain Control: R^2^ = 0.497 ± 0.15; paired t-test: p = 0.0007, n = 19 cells). These results demonstrate that gain control mechanisms are necessary to predict responses to a succession of two natural images, especially in W3 cells.

We then tested the ability of this model to reproduce experimental LSTAs by computing the gradient of the model output relative to the input probe (see Methods)^22^. The model successfully predicted experimentally observed LSTAs (Fig. 2E, F). We estimated prediction quality via correlation between predicted and experimental LSTAs (see Methods). The Gain Control CNN significantly outperformed a LN model baseline (approximated by the classical STA) in W3 cells, achieving a correlation of 0.48 compared to 0.36 for the control (Fig. 2G; paired t-test, p < 0.001, n = 180 LSTAs). While less pronounced, this improvement extended to all cell types, with correlations of 0.56 versus 0.53 (paired t-test, p < 0.001, n = 1746 LSTAs).

The Gain Control CNN thus demonstrates a robust capacity to generalize and predict how a previously unseen adaptor will modulate not only a cell’s response but also its spatial selectivity and polarity. This suggests that retinal ganglion cell selectivity in natural contexts relies on a history dependent gain control that is effectively captured by our biologically inspired architecture.

### Gradient geometry reveals a selectivity for homogeneous regions

How does the gain control mechanism impact ganglion cell selectivity to natural scenes? To answer this, we examined the relationship between images and gradients of the cell’s functional response (i.e. LSTA), following an approach similar to Goldin et al^16^.

We used the Gain Control CNN model to generate predicted gradients for all the natural probes (grey adaptor) in the training set for each cell (Fig. 3B). We then applied principal component analysis to these predicted gradients for each cell (Fig. 3A), constructing a low-dimensional representation of stimulus space that captures most of the variability within gradients.The first two principal components (Fig. 3A) accounted for a large fraction of the variance of gradients across cells (81 ± 17% across all cells). We then projected both the images and their associated gradients into this two-dimensional space, representing each image as a point and the corresponding gradient as an arrow (Fig. 3C; see Methods). Each arrow indicates the direction of change in the stimulus space that would most effectively increase the firing rate of the cell locally.

**Fig. 3.**
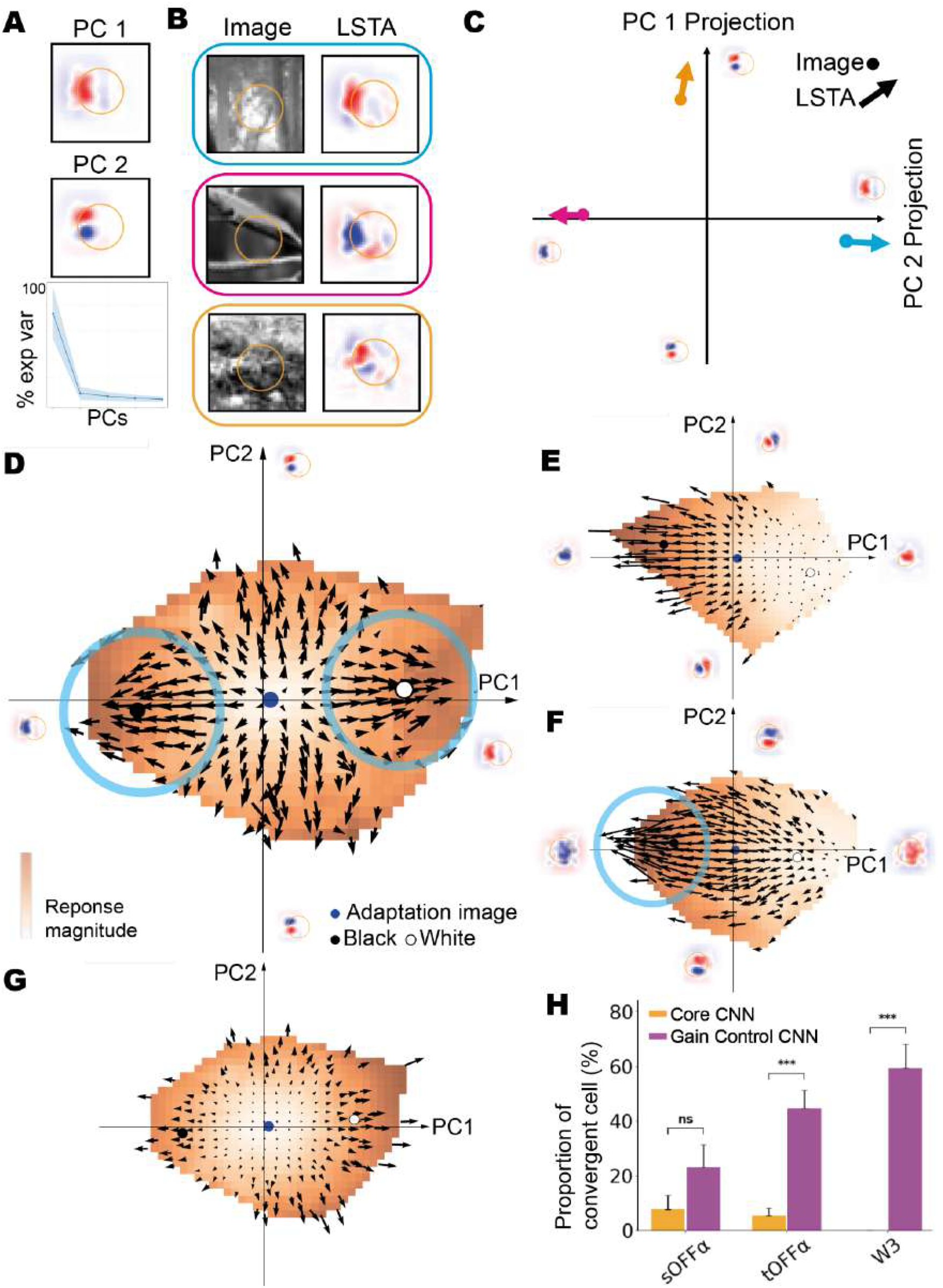
Gradient geometry reveals a selectivity for homogenous regions. **A**. For each cell, we predicted LSTAs for 3190 natural images (after grey adaptation) with the CNN model and performed a PCA on them. Top and middle row: the first two PCA components for an example W3 ganglion cell. Bottom row: variance of the LSTA ensemble explained by the first five principal components, averaged across all modelled cells. **B. C**. Representation of image-LSTA pairs as points and arrows in the two dimensional PCA space representation for the stimulus space. In B. three image-LSTA pairs are color coded into point-arrow pairs. In C. each arrow origin is located on the projection of one image in the PCA space and each arrow indicates the projection of one image once superimposed with its LSTA. **D**. Gradient field for an example W3 cell. The color overlay indicates the average firing rate to images in that part of stimulus space. Regions of convergence of the gradient field are highlighted by the blue circles. **E**. Gradient field for an example sustained-OFFα. **F**. Gradient field for an example transient-OFFα. **G**. Same as D but using the core CNN. **H**. Frequency of convergence across common cell types. Gain control significantly increased gradient field convergence in transient-OFFα (p < 1×10^−4^, n = 56 cells) and W3 cells (p < 1×10^−4^, n = 32 cells), but not in sustained-OFFα cells (p = 0.13, n = 26 cells; χ^2^ test throughout). Data are presented as mean and SEM.

**Schematic A.**
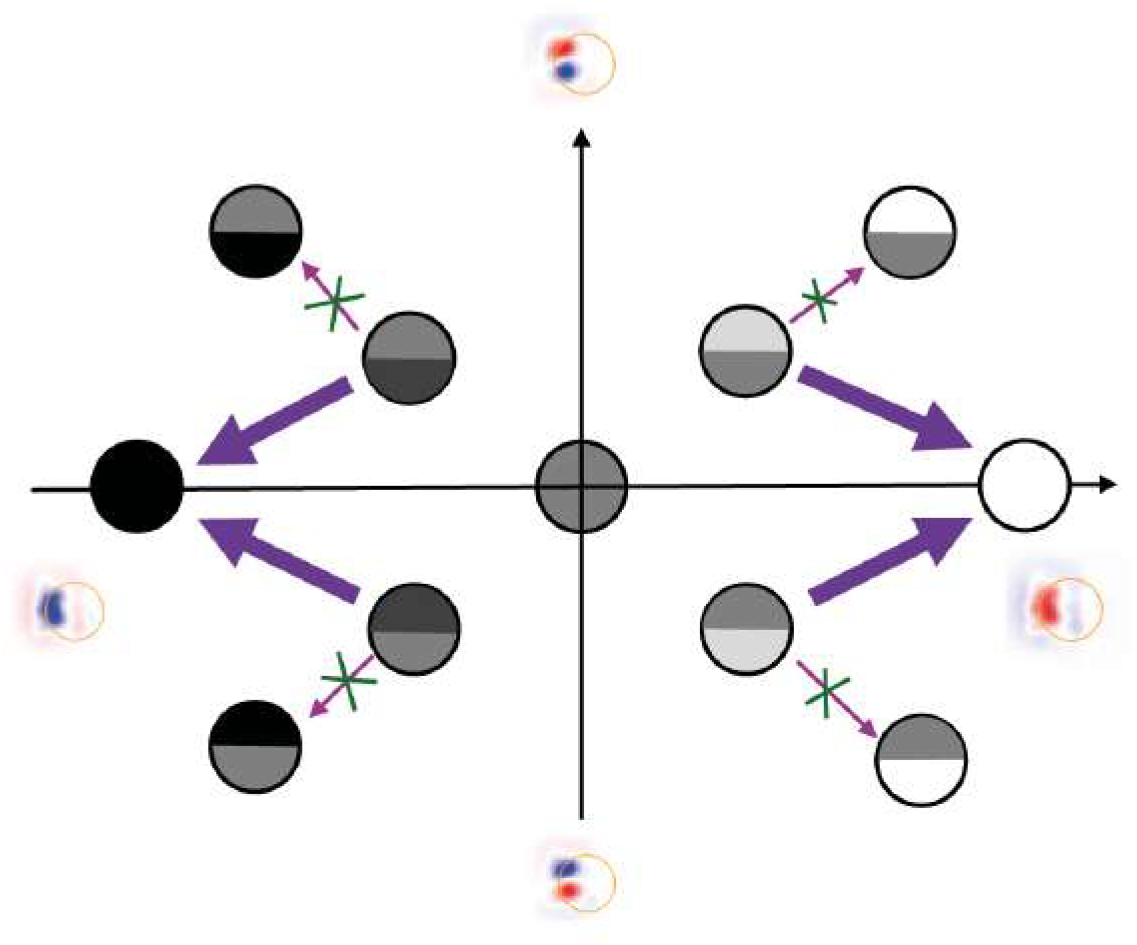
Gradient fields convergence points to homogeneity. Circles show example stimuli in the 2D PC space of a W3 cell; arrows show the gradient (LSTA) predicted by the Gain Control CNN, i.e. the direction that most increases firing. Small red-crossed arrows show the Core CNN’s gradient for comparison. PC1 (x-axis) is a blob-like filter capturing mean luminance; PC2 (y-axis) is an edge-like filter capturing top/bottom contrast polarity. Take a globally dark edge pattern, darker at the bottom (top-left of the space). The Gain Control CNN’s gradient points toward extreme negative PC1: a uniform black image. This gradient is negatively aligned with PC2, i.e. toward less contrast, not more. The mirror case (bright-on-top edge, top-right) points toward uniform white, again suppressing local contrast. The Core CNN’s gradient does the opposite in both cases: it points outward, increasing edge contrast rather than suppressing it.

We observed that for W3 cells, the gradient arrows across stimulus space converge toward two preferred regions of stimuli space (regions of convergence are highlighted with a blue circle in Fig. 3D and F). These regions lie along the PC1 axis, corresponding to near-uniform black or near-uniform white images within the receptive field (see Supp. Fig. 1 and Methods). Consequently, W3 cells demonstrate a clear preference for homogeneous stimulation in their receptive field. For example, if an image contains an edge where one half is darker, the optimal way to increase the cell response is to darken the remaining half rather than increasing existing spatial contrast (see Schematic A). Similarly,in the transient-OFFα type, convergence also happened but only in the dark portion of stimulus space since those cells did not display ON LSTAs for any natural images

For comparison, sustained-OFFα cells presented arrows pointing consistently in one direction throughout the space, indicating that their gradient shape and polarity remained relatively constant regardless of the image they were measured around (Fig. 3E). For these cells, the preferred change in the stimulus is always the same (a decrease in luminance in the receptive field for the example in Fig. 3E).

Interestingly, gradient fields of W3 and transient-OFFα predicted by the Core CNN did not display any convergence (Fig. 3G), suggesting that gain control is indeed required for this property. To determine which cell types show this convergent pattern, we estimated the local divergence of the gradient fields predicted by both models and tested for each cell if it was significantly negative (see Methods). This confirms that convergence was much more likely to happen in the transient-OFFα and W3 cell types (Fig. 3H, see Methods). This convergence was always toward an homogeneous area of stimulus space: convergence points are close to the PC1 axis and PC1 is largely homogeneous in the receptive field (see Supp. Fig. 1). The preference for homogeneous stimuli was thus specific to these ganglion cell types and enabled by gain control mechanisms.

### Gain control enables specific cell types to detect homogeneous objects situated within cluttered natural background

Our analysis of the model predicted gradient fields suggests that W3 cells are selective to homogeneous stimuli inside their receptive fields. We hypothesize that this selectivity to homogeneity enables these cells to detect spatially uniform objects situated within cluttered backgrounds. To test this, we designed a two-frame stimulus: a naturalistic background presented alone, followed by a homogeneous eagle silhouette overlaid in the same scene.

For each cell type, we predicted the response of a mosaic of cells of such type by spatially convolving the gain control model fitted for one cell of the type across the image sequence (see Methods). As hypothesized, the response to the uniform object (the eagle) “popped out” more clearly in the W3 mosaics that exhibit homogeneity selectivity (Fig. 4B). In contrast, sustained-OFFα cells responded non-selectively to both the object and background, failing to set the eagle apart from the background.

**Fig. 4.**
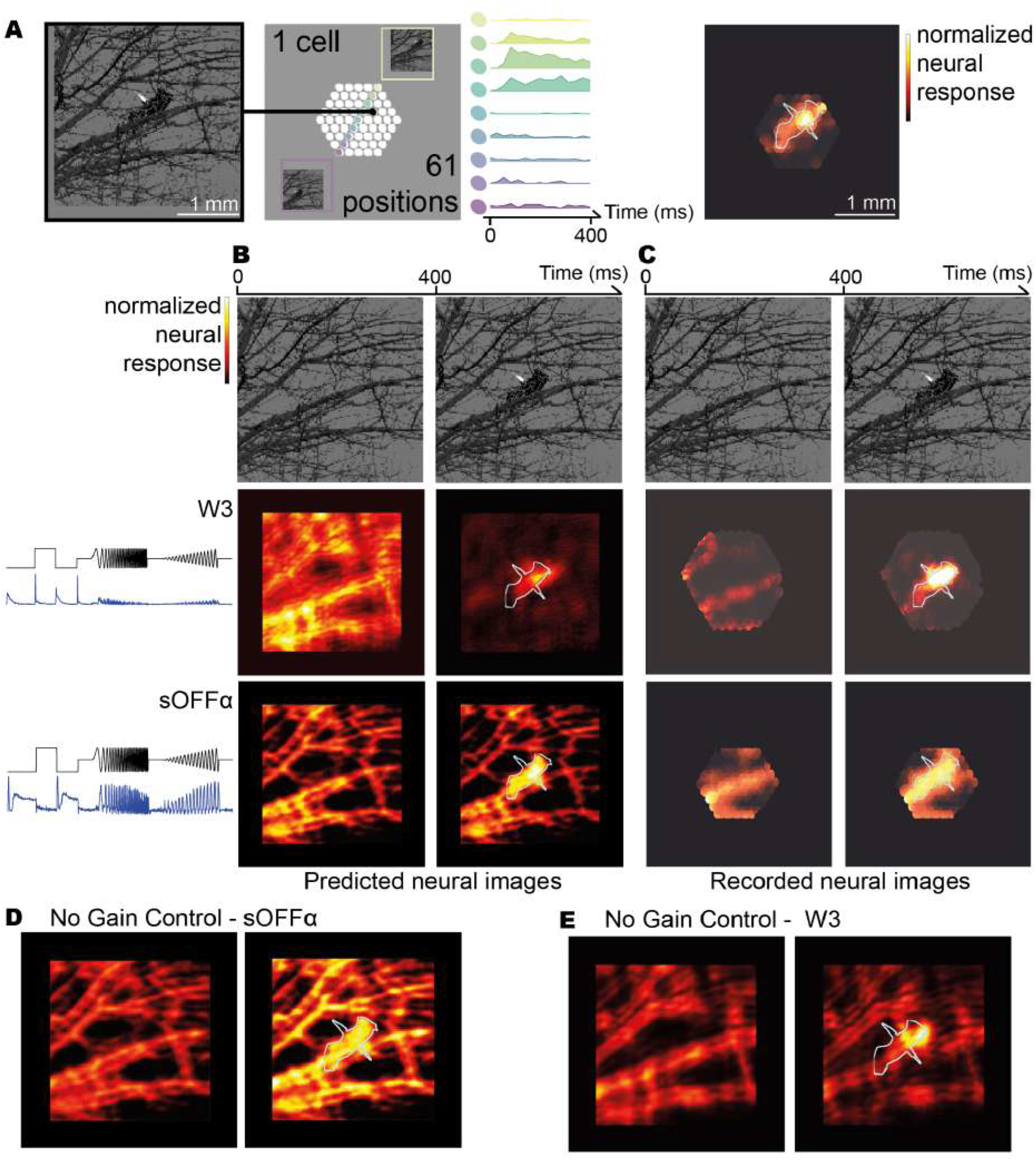
Gain control enables specific cell types to detect homogeneous objects situated within cluttered natural background. **A**. Schematic of the experimental recording of neural images. The same sequence was presented with 61 different spatial shifts (left panels), and we recorded the activity of each retinal ganglion cell at each position (30 repetitions per shift, middle panel). To construct the neural image, we color the receptive field at each position depending on the firing rate of the cells when the movie was shown at this position (right panel, see Methods). The color scale represents the mean firing rate at each pixel, normalized individually for each sequence. The contour overlay indicates the position of the object in the stimulation. **B**. Gain Control CNN prediction of the neural images for a sequence made of a background followed by the same background with an object in it. For each type, the average response to the ‘chirp’ stimulus average across the population is plotted to the left (x axis : time, top y axis : luminance level of the visual stimulation, bottom y axis : average firing rate, see Methods). **C**. Recorded neural images for the same sequence as B. (sOFFα : n = 3 cells, W3 : n = 15). Those population neural maps are the average of the neural maps of all cells in the population (see Methods). **D, E**. Same as B but without the gain control layer (Core CNN).

To verify this experimentally, we reconstructed high resolution neural images of cell mosaics from ganglion cell recordings. Because a single recording typically does not capture a perfectly tiled mosaic of one cell type, we used a spatial displacement protocol (Fig. 4A). We recorded each cell’s responses to the same movie at 61 spatial offsets, mapping firing rates to their corresponding positions. We then averaged these mappings across cells of the same type, accounting for their relative receptive field positions. Experimental data confirmed that W3 cells exhibited greater selectivity for the appearing object compared to sustained-OFF α cells (Fig. 4D).

To isolate the contribution of gain control, we performed an in-silico ablation: removing the gain control layer from the model (effectively using the Core CNN) significantly reduced the ability of W3 cells to suppress background noise and isolate the target (Fig. 4D, E). This further confirms that the gain control mechanism is necessary for homogeneity selectivity in those cell types.

In W3 cells, LSTA convergence was observed for both dark and bright stimulus regions (Fig. 3I), suggesting these cells detect homogeneity regardless of contrast polarity. This further tells W3 apart from other homogeneity detectors such as transient-OFFα that do not respond to bright stimuli^23^. We therefore predicted and experimentally confirmed that W3 cells selectively respond to a bright object embedded in the same cluttered background (Fig. 5A, B), further demonstrating that gain control supports homogeneity detection across both contrast polarities.

**Fig. 5.**
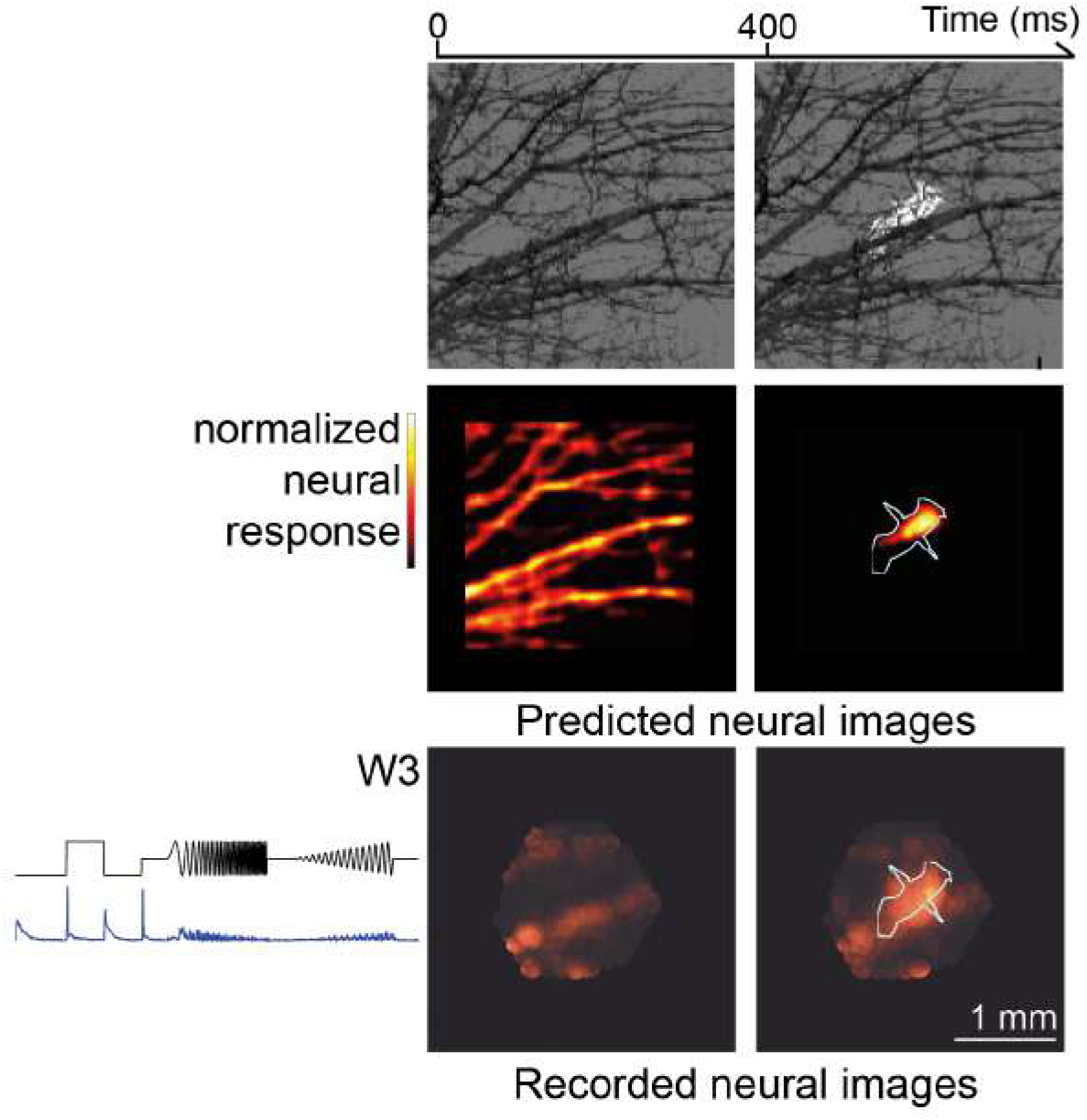
W3 cells can detect objects regardless of the object luminance polarity. **A**. Gain Control CNN predictions of the neural images for the same sequence as Fig. 4 with reversed contrast of the object. The color scale represents the mean firing rate of the cell mosaic at each pixel, normalized individually for each sequence. The contour overlay indicates the position of the object in the stimulation. **B**. Recorded neural Images for the same sequence as A (W3 : n = 14 cells). The color scale represents the mean firing rate of the cell mosaic at each pixel, normalized individually for each sequence. The contour overlay indicates the position of the object in the stimulation. Mean responses across cells to the chirp stimulus are presented on the left (see Methods).

Together, these results show that gain control does more than normalize responses. Even at the earliest stage of visual processing, it actively reshapes retinal feature selectivity to support robust object detection in naturalistic conditions.

## Discussion

We have shown that fast adaptive mechanisms, that are crucial to predict how ganglion cells generalize to new input statistics, have additional roles. They substantially reshape the selectivity of some ganglion cell types during natural stimulation, to make them primarily sensitive to homogeneous stimuli inside their receptive fields. Overall, this reframes fast gain control as an enabler for new feature detection, rather than just a corrective mechanism for keeping neurons in their operating range. This also suggests that biological gain control mechanisms contribute far more to visual encoding than simpler normalizations, such as batch normalization, typically found in deep neural networks.

Encoding the ever changing visual world is a major computational challenge. Biological systems address this through saccade-and-fixate gaze strategies to actively slice the continuous visual scene into a sequence of discrete fixation^15,24^. The rapid image transitions generated by these gaze shifts trigger short-term adaptation, changing the feature selectivity of retinal ganglion cells within each individual fixation^6^. Our results show that gain control allows specific types of ganglion cells in the retina to adapt to the cluttered background of natural scenes, and extract selectively large homogeneous objects. The combination of saccades and non-linear processing in the retina make specific features pop out in the retinal representation, something that could not be done by a passive relay of information.

### Models of adaptation and generalization

A central challenge in building models of sensory processing is generalization. A model that fits responses within a single stimulus distribution may do so by learning statistical regularities specific to that distribution, rather than capturing the underlying biological mechanism. Consequently, traditional receptive field models optimized for artificial stimuli fail dramatically when predicting responses to natural scenes, both in V1^25–27^ and in the retina^28,29^, and vice versa^30^. Here we show that a model which generalizes across complex visual statistics is necessary to uncover the function of specific neuron types. This necessity follows directly from how gain control operates. In our model, local gain control acts divisively on each subunit’s output, immediately after the subunit’s own nonlinearity. The stronger a subunit is activated, the more its output is suppressed. At moderate activation levels this suppression is weak, so grey-adapted responses are barely affected; at high activation it becomes strong, suppressing that subunit’s contribution. A sparse stimulus drives a few subunits strongly, gain control suppresses their responses heavily, and the reduced sum often fails to clear the final threshold. Conversely, a homogeneous stimulus drives many subunits moderately, so gain control barely suppresses them, and their sum more easily clears it. A model trained on grey adaptation alone could instead mimic convergence of the gradient field (Fig. 3D) with a high, fixed, non-adaptive final threshold, indistinguishable from real gain control unless tested across other adaptive contexts. Only by testing generalization across multiple adaptive contexts can we distinguish a model that has learned the true adaptive mechanism from one that has merely learned a distribution-specific shortcut. Solving the generalization problem is thus not a mere engineering exercise, but is required to correctly attribute function to specific neuronal types. A mechanistically inaccurate model can otherwise misidentify the true computation entirely.

Our model combines two distinct strategies to achieve robust generalization: some elements embed fixed statistics, while others provide dynamic mechanisms to adapt when those statistics change. First, when the statistical properties of the visual data are stable, a model can simply learn those average properties from the training set. For instance, Batch Normalization (BN) at the subunit and readout levels normalizes activations based on the mean and variance of the data. By tracking these running statistics during training, BN effectively removes baseline luminance and contrast variations. This serves as a computational surrogate for a form of slow adaptation^31^. Because luminance and broad contrast remain constant between our adaptors and probes, this fixed approach is a good approximation. However, batch normalization differs strongly from gain control. When the visual stimuli vary dynamically (here when the adaptor changes), learning the average is no longer sufficient. This is where an explicit biological mechanism becomes necessary. Our model handles this variation through a dynamic gain control layer. Rather than absorbing adaptation into static parameters, this layer encodes the underlying computational principle directly, enabling the model to generalize to entirely unseen stimulus histories. As a consequence, while batch normalization only models a normalization of the input, gain control can be used to make cells selective to specific features, like homogeneity.

It is instructive to compare our approach to solve the generalization problem to the strategy of foundation models^32^. Foundation models aim at massively increasing the training set to improve generalization performance, which allows to increase the number of parameters of the model. They thus differ from our strategy in two ways. First, we aim at minimizing the number of parameters that need to be added to achieve generalization. In our case, the gain control component had only one free parameter. Second, in most cases so far, the test for generalization has been to predict responses to simpler, classical stimuli. In our case, we switched the adaptor from grey to a natural image, a manipulation designed to generalize to a substantially more complex stimulus.

More broadly, scaling up data reflects a bet that, given enough diverse training data, a model will eventually discover the right biological computations on its own, even though it relies on a generic architecture that carries its own biases (e.g., recurrent layers, normalization layers). An alternative is to impose the relevant structural bias directly, from known biology. Incorporating biological mechanisms as inductive biases has proven effective at multiple stages of the visual hierarchy: photoreceptor-level gain control enables generalization across luminance conditions^13,33^, and various architectural constraints improve generalization across stimulus statistics in both the retina^30^ and visual cortex^27,34,35^. A natural extension of this logic is that combining multiple such constraints, each targeting a different level of the visual hierarchy, could yield ‘hybrid’ models that generalize even more broadly^36^. Our work shows that this ability to generalize is key to building neural models that fully capture the function of visual cells in the early visual system.

### Circuit mechanisms of adaptive homogeneity selectivity

Several retinal circuits could plausibly implement gain control after the first layer of our model. Specifically in W3 cells, the local spatial scale of gain control in our model points toward circuit mechanisms operating at intermediate levels of the retinal circuitry. One of them is short-term synaptic depression at bipolar cell terminals that limits excitatory drive during prolonged stimulation^37–39^. Another source of adaptation is feedforward inhibition from amacrine cells that shapes the ON/OFF balance and contributes to distinct forms of contrast adaptation within these local circuits^40–43^. Glycinergic narrow-field amacrine cells would appear as a prime candidate^44^.

Another candidate mechanism is photoreceptor-level adaptation that would occur at the finest spatial scale and may also propagate downstream^33^. However, adaptation happening purely at the photoreceptor level is unlikely to produce distinct changes in feature selectivity between different cell types that receive light signals from a similar pool of photoreceptors. We tested the ability of a biophysical photoreceptor layer as a front end to our Core CNN to predict responses to adaptor-probe sequences. This model did not capture cell responses as well as the Gain Control CNN, even after fine-tuning the photoreceptor layer on responses with different adaptors (Supp. Fig. 3B, see Methods). This performance difference is more pronounced in adaptive cell types (W3 and tOFFα) than non-adaptive cell types (sOFFα). This difference between cell types argues against an explanation at the level of photoreceptors, which would affect all cell types similarly.

Global adaptive mechanisms, often intrinsic to the ganglion cell like slow inactivation of voltage-gated sodium channels or afterhyperpolarization currents^45–48^, act after spatial integration. They are unlikely to explain our results alone. A global gain mechanism can’t explain the sign inversion of the LSTA, because a purely divisive rescaling downstream of spatial integration only scales the response. It can’t change the LSTA’s shape or flip its ON/OFF polarity. To confirm this, we compared placing the gain control either after individual subunits to model local gain control over a bipolar cell-sized spatial pool, or after the full spatial integration stage to model global gain control over the entire receptive field. While for most types the location of the gain control did not matter, the performance of the W3 model was significantly degraded when using the global gain control (see Supp. Fig. 2). This aligns with results from Garvert and Gollisch49 demonstrating that adaptation is mostly global in the receptive field, in most but not all retinal ganglion cells.

For comparison, a recent study by Chen et al.^45^ uncovered the circuitry driving the homogeneous selectivity of tOFFα cells. They demonstrated that this tuning emerges from a combination of spatially local inhibition from AII amacrine cells and strong short-term synaptic plasticity at the bipolar terminal. Their characterization was conducted at significantly lower light levels than the current study (~100 R*/cone/s vs. ~10^5^ R*/cone/s), and their stimulation targeted only the center of the cell’s receptive field, leaving open whether the same circuit mechanisms operate across both studies. Notably, they report that tOFFα cells lose their homogeneity selectivity at higher light levels. This could explain why tOFFα cells demonstrated weaker homogeneity selectivity than W3 cells in our experiments at photopic light levels (see Fig. 3H).

### Functions of the W3 cell types

The most adaptive cell type in our analysis, the W3 type, may correspond to more than one cell type^46,47^. It is thus unclear whether all those subtypes are selective for homogeneity. Our W3 recordings are most consistent with the HD2 subtype as described in Jacoby et al.^48^. HD2 cells have small receptive field centers with strong surround suppression. Postsynaptic inhibition truncates their spike responses, aligning with the transience we observe from gain control. Their ON responses are also not fully suppressed by large stimuli, consistent with our ‘W3’ data. One potential discrepancy is that HD2 cells were reported as OFF-dominant under scotopic and photopic conditions. While W3 cells in our model are more strongly connected to OFF subunits (Supp. Fig. 3), W3 cell responses in the data appear more ON-dominated under full-field photopic stimulation (chirp responses, Fig. 4B). Experiments using genetic markers paired with functional typing could confirm the correspondence between the cells we classify as W3 and HD2. New standard stimulation to identify functional cell types in the mouse retina along the chirp stimulus could also help resolve this lack of precision.

Other W3 subtypes are also known to be powerful detectors of ecologically relevant visual features. Along with tOFFα, one of them is known to respond to looming stimuli, though in functionally distinct ways: tOFFα encodes the speed and trajectory of an approaching object, while W3 signals its onset without conveying motion information^49^. Whether the gain control mechanism we identify here also contributes to looming detection in visually cluttered environments remains an open question.

### Gain control compensates for global and local spatial contrast in W3 cells

What is the consequence of the locality of this gain control mechanism ? While we investigated the feature selectivity enabled by gain control, we have not yet established how the adaptor itself (the recent stimulus history) shapes this selectivity. Specifically, it is unclear whether adaptation driven by global, full-field features and adaptation driven by local, spatially restricted features would produce distinct changes in selectivity, and through what mechanisms these two components might interact. We used the gain control model to explore this question. When adapted to a bright natural scene, W3 cells appear to lose their ON selectivity and behave like pure OFF cells, with gradient field arrows converging strongly in the negative luminance region of stimulus space (Fig. 6C). This would enhance their selectivity for dark homogeneous regions, such as the dark eagle in Fig. 4, making them indiscernible from transient-OFFα cells in this temporal context (Fig. 3F). This may also explain the polarity inversion of some LSTAs observed experimentally (Fig. 1E, 6G). Symmetrically, when adapted with dark natural scenes, W3 cells appeared as pure ON cells.

**Fig. 6.**
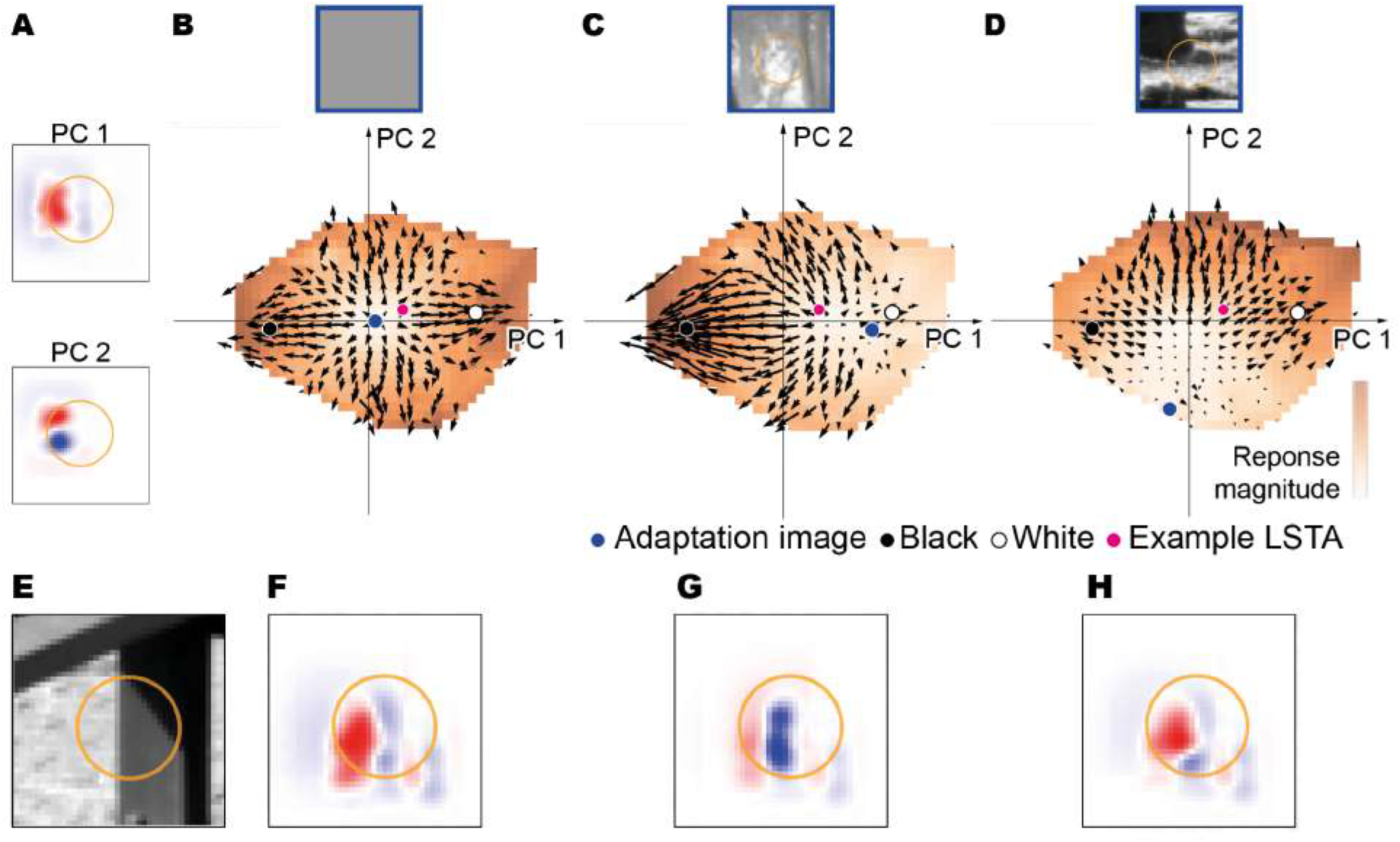
Gain control compensates for global and local spatial contrast in W3 cells. **A**. The first two PCA components for an example W3 ganglion cell. (same cell as Fig. 3) **B**. Gradient field for which the adaptor is the indicated natural image (indicated in the gradient field by the blue dot). All LSTAs are predicted using the Gain Control CNN model. **C. D**. Same as B for different adaptors. **E**. Example Natural Image (indicated in the gradient field by the pink dot) **F**. Predicted LSTA to the image in E after the adaptation as in B **G**. Same as F but with adaptation as in C **H**. Same as F but with adaptation as in D

**Schematic B.**
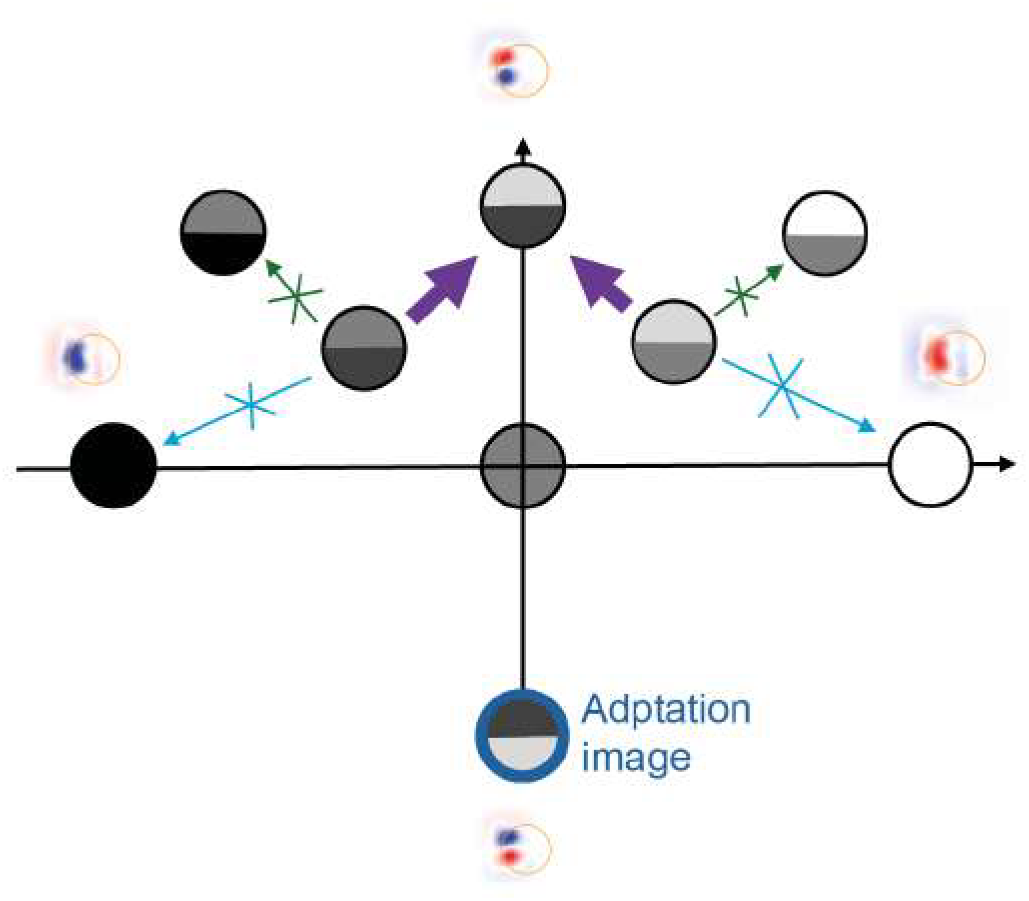
Effect of the adaptor on the gradient field. Purple arrows show the gradient (LSTA) predicted by the Gain Control CNN for the W3 cell after adaptation to the image circled in blue. Thin green arrows show the Core CNN’s gradient for comparison. Thin blue arrows show the Gain Control CNN’s gradient after adaptation to a grey (neutral) image. See the Fig. 3D schematic for further details on the axes and interpretation.

A second effect emerges when the adaptor matches edge-like features (PC2, y-axis). Here, gain control appears to act as a locally compensatory mechanism: the cell preferentially responds to stimuli opposite to it (Fig. 6D, Schematic B). This could explain the spatial shift in LSTA centers observed experimentally (Fig. 1D, 6H). While this might look like a total shift in feature preference from spatial uniformity to contrasted edges, it could actually demonstrate that W3 cells dynamically compensate for the adaptor’s spatial structure. By leveraging gain control to prioritize stimuli that restore uniformity to the visual input stream, these cells continuously reshape their feature selectivity based on recent visual history.

## Acknowledgments

We would like to thank Matthew Chalk for his help with modelling and writing and all the members of the Marre lab for helpful discussions. This work was supported by the ERC Consolidator grant DEEPRETINA (101045253) (O.M.), ANR grant Chaire Industrielle MyopiaMaster ANR-22-CHIN-0006 (O.M.), ANR grant ANR18-CE37-0011–DECORE (O.M.), ANR grant ANR-20-CE37-001804–Shooting Star (O.M.), ANR grant ANR-22-CE37-0033 NUTRIACT (O. M.), ANR grant project ANR-22-CE37-0016-01 PerBaCo (O.M.). ANR grant project RetNet4EC (O.M.), AVIESAN-UNADEV (O.M.), Retina France (O.M.), Programme Investissements d’Avenir IHUFOReSIGHT 497 (ANR-18-IAHU-01) (O.M.). S.V. was supported by the European Union’s Horizon 2020 research and innovation program under the Marie Skłodowska-Curie grant agreement No 861423. O.M.’s lab is part of the DIM C-BRAINS, funded by the Conseil Régional d’Ile-de-France.

## Author contributions

Conceptualization: BL, SV, OM; Methodology: BL, DV, RB, SV, OM; Software: BL, SV; Formal analysis: BL; Investigation: BL, DV, RB, SV; Visualization: BL; Supervision: SV, OM; Writing— original draft: BL, OM; Writing—review & editing: BL, SV, OM.

## Declaration of generative AI and AI-assisted technologies in the manuscript preparation process

During the preparation of this work, the author(s) used Claude (Anthropic) and Claude Code for assistance with writing, data analysis, and figure preparation. The author(s) reviewed and edited the output as needed and take full responsibility for the content of the published article.

## Code and Data availability

Code and data used for modelling and in silico model analysis will be available in OpenRetina :https://github.com/open-retina/open-retina

Code for the analysis of MEA data, cell typing, and visualization of neural images will be made available at :

https://github.com/retinal-information-processing-lab/Standard_analysis_pipeline

More data (neural image and LSTA) can be made available upon request to the authors.

## Methods

### Electrophysiological recordings

Electrophysiological data were recorded from isolated retinas from 5 C57BL6J mice of 10 to 23 weeks (median of 10). The animals were housed in enriched cages with ad libitum food, and watering. The experiment was performed in accordance with institutional animal care standards of Sorbonne Université. After euthanizing the animal, the eye was enucleated and transferred rapidly into oxygenated Ringer medium (for 2L: NaCl 125 mmol/L (14.61 g), KCl 2.5 mmol/L (0.373 g), MgCl2·6H2O 1 mmol/L (0.407 g), NaHCO3 26 mmol/L (4.369 g), CaCl2 1 mmol/L (0.294 g), L-glutamine 0.43 mmol/L (0.126 g), NaH2PO4 1.25 mmol/L (0.300 g), Glucose 20 mmol/L (7.206 g)). Dissection was made under dim light conditions as described previously^50,51^. We mounted a piece of retina onto a membrane, and then lowered it with the ganglion cell side against a 252-channel multi-electrode array (MEA) whose electrodes were spaced by 30 μm. During dissection and recordings, the tissue was perfused with oxygenated Ringer medium and a peristaltic perfusion system with 2 independent pumps: PPS2 (Multichannel Systems GmbH). Mice retinas were kept at 35–37 degrees. The data sampling rate was 20 kHz. The raw signal was acquired through MC_Rack Multi-channel Systems software 4.6.2, it was highpass filtered at 100 Hz, and the spikes were isolated using SpyKING CIRCUS 1.0.628. Subsequent data analysis was done with custom-made Python codes.

For LSTA and modelling we extracted the activity of a total of 1151 neurons from 4 experiments. We kept cells with a low number of refractory period violations (< 0.5% for all experiments, 2 ms refractory period, average RPV of conserved cells : 0.084 ± 0.12%) and whose template waveform could be well distinguished from the template waveforms of other cells. These constraints ensured a good quality of the reconstructed spike trains. For the Neural Image experiment, we extracted the activity of 203 cells with a low number of refractory period violations (< 0.5% for all experiments, 2 ms refractory period, average RPV of conserved cells : 0.23 ± 0.22 %).

### Visual stimulation

A white mounted LED (MCWHLP1, Thorlabs Inc./X-Cite exact, excelitas) was used as a light source, and the stimuli were displayed using a Digital Mirror Device (DLP9500, Texas Instruments / DLP DAD2000, Texas Instruments, pixel size projected on the retina : 3.5/2.6 μm) and focused on the photoreceptors using standard optics and an inverted microscope (Nikon). The mean light levels used for all stimuli are in the range of photopic vision: 1×10^6^-3.4×10^6^, 5.5×10^3^-1.3×10^4^, 5×10^5^-1.46×10^6^ isomerisations / (photoreceptor. s) for rods, S cones and M cones respectively.

#### Checkerboard stimuli

We displayed a random binary checkerboard during 40 min to 1 h at 30 Hz to map the receptive fields of ganglion cells. Check size was 42 μm for mice. A three dimensional STA (x, y and time) was sampled using 21 time samples. The spatial STA presented across all the figures was obtained as the 2 dimensional spatial slice at the maximum value after smoothing. The temporal STA is the one dimensional time slice at that same value. A double Gaussian fit was performed on the resulting spatial STA, and the ellipse corresponding to a 1.5σ contour of the fit was plotted for all the figures.

#### Shifting White Noise stimuli

To estimate receptive fields with higher spatial accuracy, we used a shifted white noise (SWN) paradigm^62^ instead of checkerboards. The stimulus was a sequence of binary checkerboards (168 µm checks, luminance randomly set to 0 or 1), with the whole grid randomly shifted horizontally and vertically by steps of 21 µm at each frame. Spatial STAs were computed from these responses and then decorrelated, by regularizing the stimulus covariance matrix and solving the resulting linear system, to remove correlations introduced by the overlapping shifts. Final receptive field profiles were obtained by fitting the decorrelated STAs with a double Gaussian.

#### Natural image stimuli

We used the Open Access Van Hateren Natural Image Dataset^14^, which consists of 4212 monochromatic and calibrated images taken in various natural environments. The calibration ensures a strictly linear relationship between scene luminance and pixel value. To avoid that the retinal system adapts to different ranges of light intensities encountered in different environments, we performed a preprocessing step. First, we identified the images with a significant number of pixels above saturation, which was defined as the proportion of pixels above a given threshold (6266 for ISO 200, 12551 for ISO 400 and 25102 for ISO 800). If the saturation level was above 2%, the image was discarded. This resulted in a total of 3190 images. Second, we cropped the central part of each image (final size: 864 × 864 px). Third, for each image, pixel values were converted to luminance with the conversion factor provided by the calibration. Fourth, the images were normalized with a custom procedure to fix the mean luminance and the root mean square (RMS) contrast: the linear scale was transformed to log scale, the distribution of pixel values centered and scaled, to finally come back to the linear scale and centered and scaled the distribution of pixel values for a second time to a final mean and standard deviation (respectively 0.5 and 0.25 for mice). Pixel values below 0 were clipped to 0, and those above 1 to 1.

#### Adaptation inducing stimuli

To create adaptation stimuli we paired a set of ‘adaptation’ images with a set of ‘probe’ images. Every different stimulus was a succession of 400 ms of adaptor and 400 ms of probe. For two experiments we used a checkerboard (checksize 126 μm) and its inverse as adaptor images, with the addition of grey as a control adaptor. For those same experiments we used ten selected natural images from the dataset described above as probes leading to 30 different pairs. The ten images were selected to contain a large variety of luminance and high contrast edges. In two other experiments we used five selected natural images (from the ten above) both as adaptor and probe. We skipped cases where the adaptation and probe would be the same and included grey as a control adaptation as well which also led to 25 recorded pairs. To assess the effect of the gain control mechanism in model performance we also used checkerboards (126 µm check size checkerboard, and its inverse) as adaptors in two additional experiments. In that case, we did record 30 PSTHs since adaptors were always different from the probes, and 9 LSTAs per cell too.

We built peri-stimulus histograms (PSTH) from the spikes evoked from all individual stimuli, using a binning of 25 ms.

#### Calibration of perturbation amplitude

Small random perturbations were applied to pixel intensities in natural images. We calibrated the perturbation amplitude as described in Goldin et al.^16^. The final perturbation amplitude chosen was 15%, where a value of 100% corresponds to the maximum intensity in the image dataset. This is slightly higher (+2.5%) than the one used in Goldin et al.^16^. These values corresponded to a change in the firing rate of approximately 1.5 Hz, in the ganglion cells that responded with a high enough firing rate to the unperturbed images.

### LSTA calculation

To record post-stimulus time histograms (PSTHs), we used 5 natural images; however, a subset of 3 images was reserved as probes to compute LSTAs. These 3 probes were presented across 4 adaptation contexts: a grey baseline and 3 natural scenes composed of the same 3 images. By omitting cases where the probe image matched the adaptation image, we recorded a total of 9 LSTAs per cell (3 for the grey baseline and 2 for each natural adaptation context).

For each reference image, we counted spikes occurring between 50 and 450 ms following each perturbed image presentation. We then calculated a spike-triggered average of the perturbation patterns, weighting each pattern by the number of spikes it evoked. The resulting LSTAs were denoised using a Gaussian process-based algorithm^52^. Because this denoising model estimates the pixel-wise uncertainty of its gradient prediction, it provided a quality selection criterion to determine whether a predicted LSTA was significantly different from zero. To determine whether a predicted LSTA was reliable enough to trust (as opposed to indistinguishable from noise), we tested it against the null hypothesis that the true LSTA is zero everywhere. Assuming the LSTA pixel values follow a multivariate Gaussian distribution with mean *μ*_*g*_ and covariance Σ_*g*_, we computed the quantity 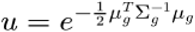 which estimates the probability of the null hypothesis (LSTA = 0) being true, smaller *u* indicates a mean vector further from zero relative to its uncertainty, i.e. a more reliable, non-null LSTA. We retained only LSTAs with *u* < 5.10^−4^.

For visualization, LSTAs were smoothed using bicubic splines, and an ellipse corresponding to the 1.5σ contour of the classical STA was overlaid as a reference in all figures.

To quantify the displacement and polarity inversion between LSTAs of the same cell responding to the same probe under different adaptation contexts, the denoised LSTAs were fitted with a 2D Gaussian envelope. Spatial distance between two LSTAs was defined as the distance between their Gaussian centers. Pairs were classified as “displaced” if this distance exceeded 0.75σ of the classical STA. This 0.75σ threshold was chosen to maximize agreement with a visual inspection baseline conducted by a single human observer on one experiment. The polarity of an LSTA was determined by the sign of its highest absolute-value pixel. Pairs were classified as having “inverting polarities” if the LSTA sign changed for the same probe image across different adaptation contexts.

To evaluate the predictive performance of our Gain Control CNN across the population of modeled cells, we used a standard Linear-Nonlinear (LN) model framework as a control. While an LN model can explain adaptive response changes via linear temporal filtering, it cannot inherently account for modulations in feature selectivity (such as displaced or polarity-inverted LSTAs). Because the LSTA of a model can be extracted via its gradient, and the gradient of a standard LN model is constant by construction^16^, we did not explicitly train an LN model. Instead, we used the cell’s receptive field estimated from a checkerboard stimulus (the spatial STA, smoothed with a Gaussian blur prior to a 2D Gaussian fit) as a proxy for the static LSTA an LN model would produce.

The LSTAs predicted by the Gain Control CNN were denoised using a Gaussian blur (σ = 63 μm) and fitted with a 2D Gaussian. Finally, both the proxy LN fits and the Gain Control CNN fits were pixel-wise correlated with the experimentally recorded LSTAs.

### Cell typing

During all experiments, we displayed a set of standardized stimuli in an attempt to type recorded functional cell types.

#### Stimulus

In addition to our perturbation protocol to detect polarity inverting cells, we applied two additional ones. 1) A full field ‘chirp’ stimulus composed of ON and OFF steps, plus varying full field frequencies and amplitudes, with luminance values ranging from 0 to 1. The stimulus is the exact same that Baden et al.^17^ used to find and classify 32 different types of ganglion cells. It was played at 50 Hz, containing 20 repetitions of 32 s length. 2) Drifting gratings (DG) moving in 8 different directions with a speed of 479.5 μm/s, at a spatial period of 959 μm (274 pixels at 3.5 μm/pixel) and at 50% Michelson contrast (0.75–0.25 luminance). Each DG lasts 10 s, preceded by 2 s of grey (0.5 luminance), the temporal period being 2 s. Therefore, each grating edge goes through the receptive fields of each ganglion cell 5 times per DG. The 8 directions were repeated 4 times in a pseudo-random manner. The stimulus profile and dynamics is identical to the one described in Yao et al.^53^ to retrieve direction selective cells. In our case we used a unique luminance value, as described in the Visual stimulation section above.

#### Typing

To cluster cells in different types, we built our analysis on the chirp and checkerboard stimulus responses, and representing each ganglion cell with a reduced representative vector. To obtain these vectors, first we constructed peri-stimulus histograms (PSTH) from the spikes evoked from the chirp stimulus, using a binning of 100 ms. Then, for each experiment, we z-scored all PSTHs and performed a PCA on them. We kept the number of components that were needed to explain 80% variance of the data (around 12 components). Second, we used the temporal profile (21 samples at 30 Hz) of the STA of each cell, obtained using the checkerboard stimulus. We z-scored it and performed a PCA, keeping the first component, which explains around 60% of the variance. This adds information about the classical STA polarity of the ganglion cells. Third, we used the area of the ellipse fitted to the classical STA, as the product value of their major and minor axis σ values. These areas were normalized from 0 to 1. In this way, we obtain a data vector of around 14 values, depending on the experiment, that describes each ganglion cell according to their response to a chirp and a checkerboard. Then, we performed an agglomerative clustering, setting the threshold value in a way that all clusters look homogeneous across PSTHs and STAs. This resulted in overclustering that produced around 50 ganglion cell groups (from around 200 cells in each experiment). In the last step, we assigned each cluster group to one of the 32 types described in Baden et al.^17^. To do this, we used the Calcium imaging data provided by the authors to match it with our data. We built from their Extended data Fig. 1, where the authors link electrophysiology and calcium imaging by means of a convolution between a Ca2+ event triggered by a single spike. We transformed our PSTHs by convolving them with a decaying exponential, in which we adjusted the temporal decay constant to maximize correlation of our cluster groups and theirs (median maximum correlation of 0.76). Cell types that present strong responses to the modulating frequencies and amplitude were assigned correctly, while other types which mostly respond to ON/OFF steps, were assigned in a second round of correlation match after excluding the former groups. Besides the correlation of the chirp traces, we confirmed the correct assigning of groups by checking that the ellipses of each type form a proper mosaic, that the spatial STAs look uniform, the similarity of their direction selectivity PSTHs (see below) and that of the spike waveforms.

For some analysis, we reported results from five non-direction-selective common types in the mouse retina (sustained-OFFα, sustained-ONα, transient-ON, transient-OFFα, W3). To identify the types mentioned in this study we used the following criteria with visual evaluation of the obtained clusters, after discarding direction selective cells.

- transient-OFFα cells have a transient response to both OFF steps and no response to the ON steps and a clear variation of the response intensity depending on the stage of the frequency.
- sustained-OFFα had a sustained response to both OFF steps and no response to the ON steps and a clear variation of the response intensity depending on the stage of the frequency.
- transient-ON cells have a transient response to both ON step and no response to the OFF steps and a clear variation of the response intensity depending on the stage of the frequency.
- sustained-ONα had a sustained response to both ON step and no response to the OFF steps and a clear variation of the response intensity depending on the stage of the frequency.
- W3 cells have a small transient response to the ON steps and a strong transient response to the OFF steps and no response to any of the frequencies.

#### Direction selectivity

We constructed PSTHs from the spikes evoked from the chirp stimulus, and calculated the mean firing rate evoked by each DG direction, and normalized it to the maximum direction for each cell (values 0 to 1). To assess selectivity, we calculated the vector sum of these normalized response vectors, which spanned values from 0 to 2, as it is usually done^17,53^.

### Adaptive Index

To evaluate how different visual contexts modified neural responsiveness, we calculated an Adaptation Ratio for each cell. This metric was designed to be agnostic to the direction of change, effectively capturing both sensitization and desensitization by measuring the absolute magnitude of the shift in peak firing relative to a baseline control.

For each experimental batch, the maximum spike count evoked during the 400 ms “probe” window following a grey-screen adaptation *MaxSpikes*_*grey*_ served as the baseline. This was compared against the maximum spike count evoked by the same probe following a naturalistic or checkerboard adaptation stimulus *MaxSpikes*_*nat*_. Values were extracted from the peri-stimulus histograms (PSTH) using 25ms temporal bins.

To ensure signal reliability and prevent the inflation of ratios by low-frequency noise, we applied a minimum activity threshold; any baseline response with a maximum lower than 5 Hz (calculated across the binning window) was excluded from the ratio calculation. The ratio for each stimulus pair was defined as:

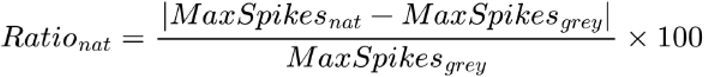

A final adaptive Index was determined by identifying the maximum adaptation ratio achieved across all tested naturalistic or checkerboard adaptation pairs for a given cell. To maintain biological plausibility and exclude potential recording artifacts, calculated ratios exceeding 200% were treated as outliers and removed from the population analysis.

### Ganglion cell modeling criteria

For the 4 mice retinas where the set of unperturbed images was presented, we selected the cells to be modeled under the following criteria: a) they did have a classical STA, b) the spiking responses across the unperturbed, repeated images were stable. To assess stability, we used the criterion of Cadena et al^27^. and discarded the neurons that showed a ratio of explainable-to-total variance smaller than 0.30. The computation of the explainable-to-total variance was done on concatenated responses from the 30 (25 for natural image adaptation) different test stimuli. We were able to model 483 cells (268 using natural images as adaptation in the test dataset, 215 using checkerboard as adaptation in the test dataset).

### Model evaluation

To evaluate the performance of the models, we used a testing set of 30 (25 for natural image adaptation) different stimuli where each stimulus has been repeated 30 times. Those 30(25) stimuli are the 30(25) adaptation pairs described above. These repetitions allowed to separate prediction error in two parts: the error due to the limitations of the model and the error due to the intrinsic noise in the response. Given n responses to the same stimulus, *y*_1_,…, *y*_*n*_, we averaged them over odd- and even-numbered trials to get two estimates of the actual mean response, *y*_0_ and *y*_e_. We defined the reliability as the correlation between these estimates, 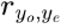. Then, given the prediction of one model, *ŷ*, we estimated the noise-corrected correlation:

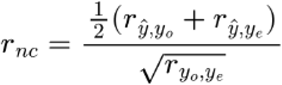

as introduced by Keshishian et al.^54^. We reported as model performance the corresponding noise-corrected r-squared, *r*_*nc*_^2^. For each cell, *r*_*nc*_^2^ was computed for each adaptor on the concatenated response to all pairs using this adaptor, then was averaged over all adaptors to report the final *r*_*nc*_^2^. Cells with *r*_*nc*_^2^ < 0.3, even after the implementation of the gain control layer, were excluded from LSTA and gradient field analysis.

We also estimated the cell reliability on the entire test dataset by computing 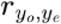 after concatenating responses to all adaptors together and we only trained models for cells with a reliability > 0.3.

### LSTA prediction

Given a model that predicts the firing rate r(t) of a neuron at time t in response to an image sequence *X*, we computed the Linear Spike-Triggered Average (LSTA). The LSTA is formally defined as the gradient of the model output with respect to the input, corresponding to the Data Jacobian matrix :

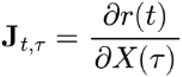

where *τ* represents the time of stimulus presentation and t represents the time of the predicted neural response. To visualize the spatial selectivity as a 2D LSTA, we computed a temporal average over a specific block of the Jacobian:

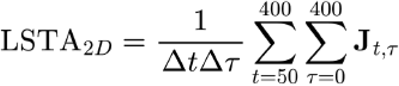

This average encompasses the period of the probe presentation (*τ* ∈ [0,400]*ms*) and the resulting neural activity (*t* ∈ [0,400]*ms*). Since all models were implemented in PyTorch^55^, we utilized automatic differentiation to compute these Jacobian blocks efficiently.

### Gradient field analysis

To understand the relationship that a ganglion cell presents between the reference images and their corresponding LSTAs, we used the CNN model to predict for each modelled cell the LSTAs for 3190 images in our dataset. A principal component analysis was made on these predicted LSTAs on a cell by cell basis to obtain a two dimensional space composed by the two first principal components (PCA1 and PCA2). We projected each image as a point in the space defined by these two components, by calculating the dot product between the image and the two components. For each image we also made the projection of their corresponding LSTA. This produced a gradient field for each cell. To ease the visualization of the gradient field, the projection space was binned and the images and LSTAs falling inside each bin were averaged to be represented by a single point (the projected image) and arrow (the corresponding LSTA).

Additionally, when plotting gradient fields resulting from non-grey adaptation (Fig. 6), we kept the PCs as obtained with the grey control adaptation.

### Gradient field convergence analysis

To assess whether or not a cell has a convergent gradient field we recorded the divergence of the binned gradient field.

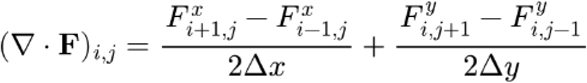

To establish a null distribution, we randomly shuffled the gradient field positions (preserving vector magnitudes and directions but randomizing spatial locations) and recomputed the divergence. A gradient field was classified as convergent if:

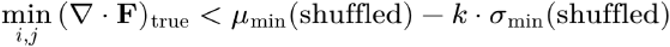

where *μ*_shuffled_ and *σ*_shuffled_ are the mean and standard deviation of the divergence values across all bins in the single shuffled gradient field, and *k* = 6 (chosen to maximize agreement with manual classification by visual inspection).

This analysis was performed separately for gradient fields computed using different adaptors (uniform grey, white, black, and the adaptation stimuli used in the test dataset: 2 checkerboards or 5 natural images). A cell was labeled as convergent if at least one of its gradient fields met the convergence criterion.

#### Convergence point and its angular coordinate

The convergence point was estimated as the occupied bin with the lowest smoothed divergence. Its angular coordinate is the angle between the vector from the origin to the bin centre and the PC1 axis, folded to [0°, 90°] so that 0° denotes a point on the PC1 axis and 90° a point on the PC2 axis. The same angle was computed for the convergence point of the shuffled field of the same cell and condition. For each cell, the angle and its shuffled counterpart were summarised as the median over all adaptation conditions that induced a convergence of the gradient field, and the two were compared across cells with a one-sided Wilcoxon signed-rank test. Because the image cloud is itself elongated along PC1, this shuffled control, rather than a uniform distribution of angles, is the appropriate null.

#### Receptive field and homogeneity of the principal components

The receptive field of each cell was defined from the two-dimensional Gaussian fitted to its checkerboard STA corresponding to a 1.5σ ellipse contour. For each principal component we computed h = ⟨PC⟩^2^ / ⟨PC^2^⟩ over the pixels inside this contour, which equals 1 for a spatially uniform component and approaches 0 for a component with balanced positive and negative lobes.

### Convolutional neural network model (CNN)

We trained one CNN model per cell in the dataset. The first layer of the CNN model was a convolutional layer with two two-dimensional convolutional kernels. The second layer was composed of one filter followed by a non-linearity. The weights of this filter were factorized in a two-dimensional spatial mask and a vector of feature weights (with one weight for each of the features extracted by the first layer) to decrease the number of model parameters, following previous work^56,27^. Finally, this one number is passed through a second non-linearity to produce the predicted response :

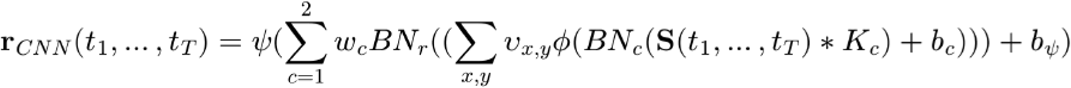

The model is composed of:

a. *First layer convolutional filter*. The filters were parametrized as the sum of a center and surround component, each factored into a spatial Gaussian and a temporal difference of Gaussians:

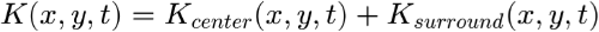

where each component is defined as the product of a spatial and a temporal kernel:

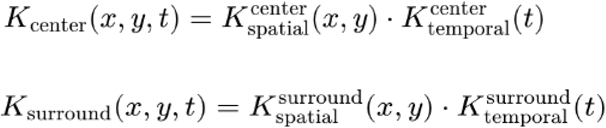

The spatial kernels are Gaussians of different widths:

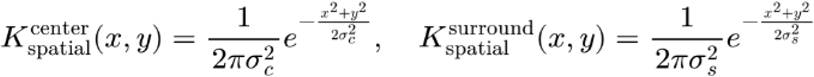

The temporal kernels are each a difference of Gaussians, allowing each component to have its own temporal dynamics:

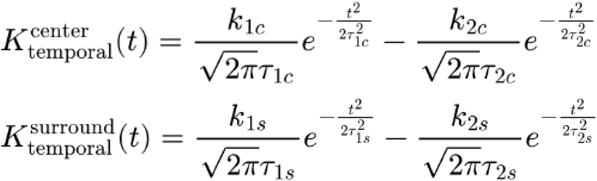

The full kernel thus reads:

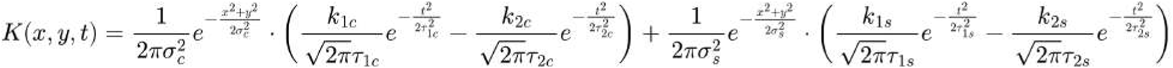

Each kernel *K* was thus parametrized by 8 learnable parameters: the spatial widths (*σ*_*c*_, *σ*_*s*_), the temporal scales 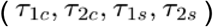, and the temporal amplitude ratios 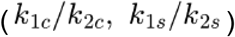 governing the biphasic shape of each temporal course. All parameters were initialized based on bipolar cell recordings^57^. All parameters were optimized through gradient descent along the rest of the model parameters (Adam optimizer). While Strauss et al.^57^ identified 14 distinct bipolar cell types, we simplified the subunit layer by averaging receptive fields separately for ON and OFF bipolar cells, resulting in a single ON subunit and a single OFF subunit. Additionally, we corrected *k*_2_ to preserve the ratio of integrated strength between center and surround when transitioning from the 1D spatial receptive fields in Strauss et al.^57^ to our 2D implementation. Since Gaussian integrals scale as k·σ in 1D but k·σ^2^ in 2D, we adjusted *k*_2_ (equivalently, *k*_1_ could have been adjusted) to maintain the original center-surround strength ratio at the model initialization. Parametrizing the subunits as differences of Gaussians conferred multiple advantages. It helps us use biological measurements to better initialise the model and reinforce mechanistic interpretability by directly linking model components to specific retinal interneurons. Finally, it reduces the number of free parameters, simplifying training and improving model stability. Because those parametrized subunits are very smooth we found the associated trained readout to be remarkably sparse, reminiscent of results from Karamanlis et al.^21^.
b. *Second layer spatial filter*. Following the subunit nonlinearities, the model performs a spatial integration of the feature maps. This stage is implemented as a spatial weighting *v*_*x,y*_ with dimensions identical to the output of the first convolutional layer. This dense spatial filter allows the model to capture the full extent of the ganglion cell’s receptive field, assigning a specific weight to each subunit’s contribution based on its spatial coordinates (*x,y*) before the final summation.
c. *Pointwise nonlinear functions. They* convert the convolutional outputs into non-negative activation values :
  th subunit nonlinearity \phi is relu

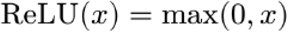
  the readout non linearity \psi is softplus

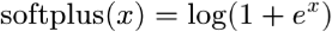
d. *Biases. (b*_*c*_, *b*_*ψ*_) prior to each nonlinearity.
e. *Readout feature weights. w*_*c*_
f. *Batch Normalization layers*. To stabilize the distribution of activations and implement a form of homeostatic gain control (as opposed to the fast dynamical gain control encoded in the gain control layer below), we integrated Batch Normalization^58^ (BN) layers immediately preceding each nonlinear activation function (*ϕ* and *ψ*). For a given input feature map *x*, the transformation is defined as:

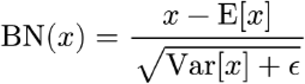

In our architecture, we purposefully omitted the learnable scaling and shifting parameters traditionally found in batch normalization layers. This ensures that the layers perform a strict standardization, forcing the model to rely on the spatial and temporal structure of the kernels rather than arbitrary parameter rescaling. At inference, *BN*_*c*_ and *BN*_*r*_ utilize the running mean and variance tracked during training. This ensures that the normalization is conditioned on the global statistics of the natural stimulus manifold, preserving the consistency of the response across varied stimuli.
g. *A Poisson noise model for training* using the Poisson negative log-likelihood loss implemented in PyTorch (PoissonNLLLoss)

#### Regularization

We used a L1 regularization on the convolutional kernel of the second layer (readout). Additionally, we treated the non-negativity of the spatial readout weights as a tunable hyperparameter.

### Model training and cross-validation

#### Model fitting

Considering the n recorded image-response pairs *X*_1_,*y*_1_, …, *X*_*n*_, *y*_*n*_, the resulting loss function is given by:

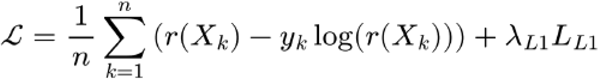

where the first term corresponds to the negative log-likelihood of the Poisson loss and where *λ*_*L*1_ and are the hyperparameters which control the importance of the L1 regularization terms. The core models were trained by minimizing a Poisson objective function using the Adam optimizer on a dataset of 2,910 natural images. Each image was preceded by a grey screen, and the target response was defined as the total spike count within a 400 ms window following probe onset. Batch sizes were varied between 16 and 256 to optimize computational throughput. We employed an initial learning rate of 0.05, managed by a “reduce on plateau” scheduler with a patience of 25 epochs. To prevent overfitting, we implemented an early stopping criterion if no improvement was observed for 50 consecutive epochs. The final parameters for each model were determined by the epoch that yielded the minimum validation loss, rather than the final training iteration.

Hyperparameters, specifically the *L*_1_ regularization strength 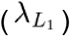 and the non-negativity constraint on the readout, were optimized independently for each neuron using a random search via the Optuna framework^59^. The optimal configuration was selected based on the minimum unregularized loss on the validation set. We utilized a 5-fold cross-validation strategy with consistent data splits across all configurations. The final model for each cell was selected from the fold yielding the lowest validation loss, ensuring robust generalization before evaluation on the held-out test set.

For each cell, images presented to the model were cropped around the center of the receptive field as estimated by the STA analysis and compressed by a factor 6 to reduce the number of pixels. Final image size in the modeling datasets were either 30 pixels of 3.5 μm (630 μm) or 40 pixels of 2.6 μm (624 μm) depending on experiments.

### Gain Control Mechanism

We implemented a gain control mechanism similar to the one described in Chen et a^22^. For each cell, two gain control layers were added (one for each subunit channel) immediately after the first-layer subunit non-linearity and before the readout layer.

In each pixel of the feature map, the new activation is :

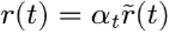

Where 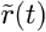 is the activation without the gain control model and

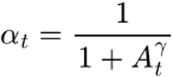

with

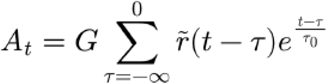

#### Optimization of the Gain Control Mechanism

This gain control layer normally introduces 3 new free parameters per subunit : *G, τ*_0_, *γ*.

The temporal constant *τ*_0_ was fixed at 400 ms, and the exponent *γ* = 2 was selected following a grid-search analysis on a representative subset of cells. Only the gain parameter *G* was optimized for each subunit (n = 2 parameters). To manage the inherent instability of *G* during optimization, we implemented a warmup scheduler, weight decay, and an early stopping strategy. All other core parameters remained frozen.

We utilized a noise-corrected correlation metric as our objective function. For each adaptation condition, the model predicted responses to the subsequent probe. We computed the correlation between the predicted and recorded responses, averaged across all probes for that condition, and normalized by the cell’s intrinsic reliability (as described in **Model Evaluation**). The final score was averaged across the 30 distinct adaptor-probe sequences (25 for natural adaptors). The final parameters for each model were determined by the epoch that yielded the minimum loss, rather than the final training iteration. As this correlation metric is scale-invariant, we performed a post-hoc rescaling of the mean predicted firing rate to match the experimental mean of the test dataset. Notably, this rescaling does not affect the reported correlation or *R*^2^ values.

Given the constraints of experimental recording time, the gain control parameters were optimized using the dataset containing adaptation stimuli (consisting of both In-Distribution and Out-of-Distribution sequences). Because the number of optimized parameters (n = 2) is negligible compared to the frozen core weights, this procedure is treated as a hyperparameter refinement rather than traditional model training. This allows us to demonstrate the performance gains attributable specifically to the gain control mechanism. Importantly, the core itself was trained on a dataset containing only In Distributions sequences (as described previously).

### Subunit polarity identification and connection strength analysis

We determined the polarity of the two subunits for each cell after training by identifying the sign of the pixel with the highest absolute value in the spatio-temporal kernels generated by the learned Difference of Gaussians (spatial and temporal) parameters. To quantify the relative contribution of ON and OFF pathways, we computed a normalized subunit connection strength.

For an individual cell *c*, let *w*_*c*,ON_ and *w*_*c*,OFF_ represent the learned linear weights of its ON and OFF subunits, respectively. We normalized these weights such that the total connection strength for each cell equals 1 (same equation for OFF):

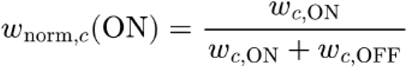

To determine the representative connectivity for a specific cell type *T*, we calculated the mean normalized weight across all *N* cells belonging to that type (same equation for OFF):

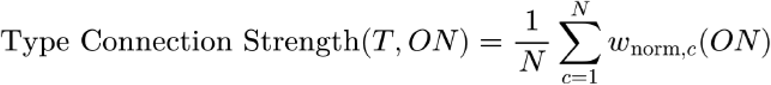

This approach ensures that the “mean” reported in our population analysis (Supp Fig. 1A) reflects the average balance of polarity inputs across cells, rather than the raw magnitude of the weights.

### Photoreceptor front end layer

To investigate whether upstream adaptation at the first stage of vision could account for the observed response dynamics, we integrated a biophysically-defined photoreceptor front-end based on the model described by Chen et al.^60^. This layer was added to the previously described Core CNN architecture.

The photoreceptor layer simulates the phototransduction cascade using a set of differential equations. Following the methodology for mouse cone parameters provided in the reference, we fixed most biophysical constants to their reported values. We designated only five key parameters as learnable to allow the model to adapt to the specific luminance dynamics of our stimuli: the gain factor (*γ*), the pigment decay rate (*σ*), the spontaneous PDE activation (*η*), the affinity of GC Ca2+ dependence (*K*_*GC*_), and the calcium extrusion rate (*β*).

The model was first initialized using the pre-trained weights of the Core CNN. During the initial training phase, the full model (Photoreceptor layer and CNN core) was trained on the grey adaptor dataset (In-Distribution). In this phase, all parameters in the model (including both the five photoreceptor parameters and the core convolutional kernels) were free to be optimized. The training strategy is the same as described for the core CNN model. Additionally, a few additional constraints were applied to keep *γ* positive and to ensure some equalities as described in Chen et al.^60^.

Following In-Distribution training, the model was fine-tuned on the test dataset containing a broader range of adaptors (Out-of-Distribution sequences). During this stage, the parameters of the Core CNN remained frozen to preserve the learned spatial and temporal features. Only the five photoreceptor parameters were permitted to change, allowing the front-end to optimize its adaptation kinetics to the complex natural and checkerboard adaptors. The optimization strategy is the same as for the gain control mechanism.

### Neural Image

#### Stimulus

We created a new stimulus by combining a context and an object picture. In the context picture, the background was defined as all pixels with values below the mean pixel value of the image. All background pixel values were set to this mean value to obtain a grey background. To create the context + object image, we first rescaled the pixel values of the object so that its darkest pixel matched the maximum value in the context foreground (100% contrast). Then, the object was placed over the background regions of the context image. This results in the object being hidden by the foreground elements of the context picture. To obtain the white object condition, we inverted the contrast of the object before display.

#### Recorded Neural Image

Experimentally, the neural image method tries to combine space and time responses of neurons in a predicted population to simulate the response of a larger population that would tile the visual field. To record neural images experimentally, we displayed the same stimulus in 61 positions forming a hexagonal grid, with a spacing of 40 μm between positions. For each cell, a neural image was constructed as follows. The position of the receptive field of the cell for each stimulus position was estimated by summing the displacement due to the stimulus shift and the position of the receptive field relative to the center of the recording. For each position, we set each set of the receptive position in the neural image to the firing rate of the cell (averaged over the entire 400 ms presentation of the stimulus). If two stimulus positions led to overlapping receptive fields, the pixel values were averaged across positions. The reliability of the neural images was computed by calculating the 2D correlation between even and odd trials (30 total trials). To obtain a population neural image, we averaged the images of all cells in the population using the following criteria: a neural image of a single cell must have reliability of at least 0.3 to be included in the average, and each pixel of the mean image must be averaged across at least two cells. When displaying these neural images, we set to zero any pixel where we did not obtain a reliable activity estimation following the above criteria and marked the boundaries of the image region where the image could be reconstructed.

#### CNN Neural Image

For the model-predicted neural image, we first selected one modeled cell of each type. We convolved each cell type model across the same stimulus, where each pixel in the predicted image corresponds to one model prediction for a cell that would be centered at that location in space. By convolving the model across the entire stimulus, we obtained a spatially complete neural image, unlike the experimental approach.

## Supplementary Figures

**Supp. Fig.1.**
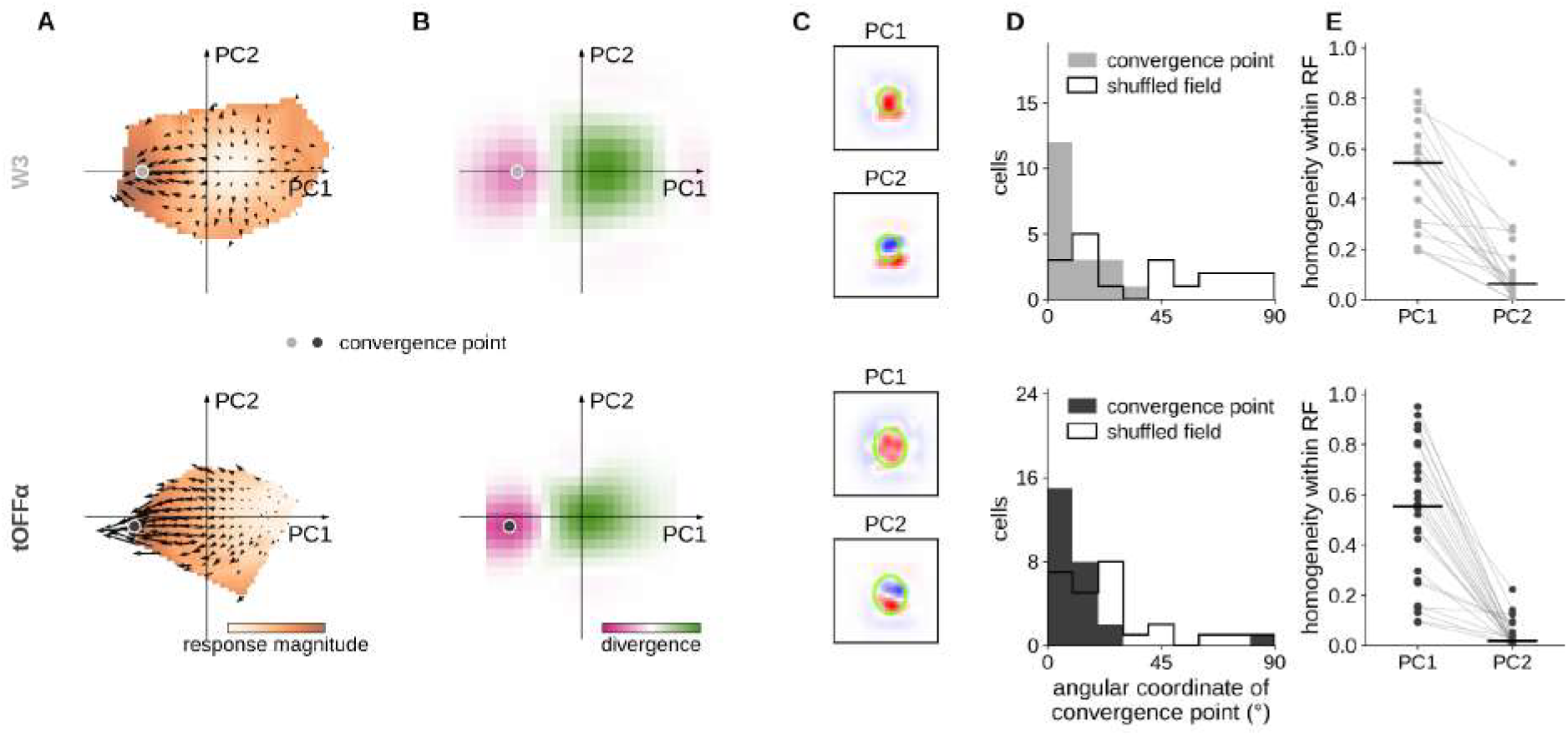
Model LSTAs converge toward a spatially homogeneous stimulus within the receptive field. **A**. Gradient field for an example W3 cell (top) and an example tOFFα cell (bottom). The color overlay indicates the average firing rate to images in that part of stimulus space. Estimated convergence points are shown as a colored dot. **B**. Smoothed divergence of the flow fields in A, using the same axes (pink: convergence, green: divergence). The convergence point is defined as the point with the lowest divergence. It lies close to the PC1 axis in both cells. **C**. PC1 and PC2 of the same cells. **D**. Angular coordinate of the convergence point (angle to the PC1 axis, projected to 0–90°) for W3 (top, n = 19 cells) and tOFFα (bottom, n = 26 cells). For each cell, we computed the median over all adaptation conditions (adaptors) that induced a convergence of the gradient field. Black outline: the same measurement on a shuffled gradient field. Convergence points are aligned with the PC1 axis (W3: median 9° vs 45° for shuffled fields, p = 0.0001; tOFFα: 6° vs 22°, p = 0.0002; one-sided Wilcoxon signed-rank test H_1_: real angle < shuffled angle). **E**. Homogeneity of PC1 and PC2 within the receptive field estimated as h = ⟨PC⟩^2^ / ⟨PC^2^⟩ over pixels in the receptive-fields (1: uniform). Each point is a cell. Median values over cells are indicated by a black bar. PC1 is more homogeneous than PC2 in every cell (W3: median 0.54 vs 0.06, p < 1×10^−4^; tOFFα: 0.55 vs 0.02, p < 1×10^−4^; one-sided Wilcoxon signed-rank test).

**Supp. Fig.2.**
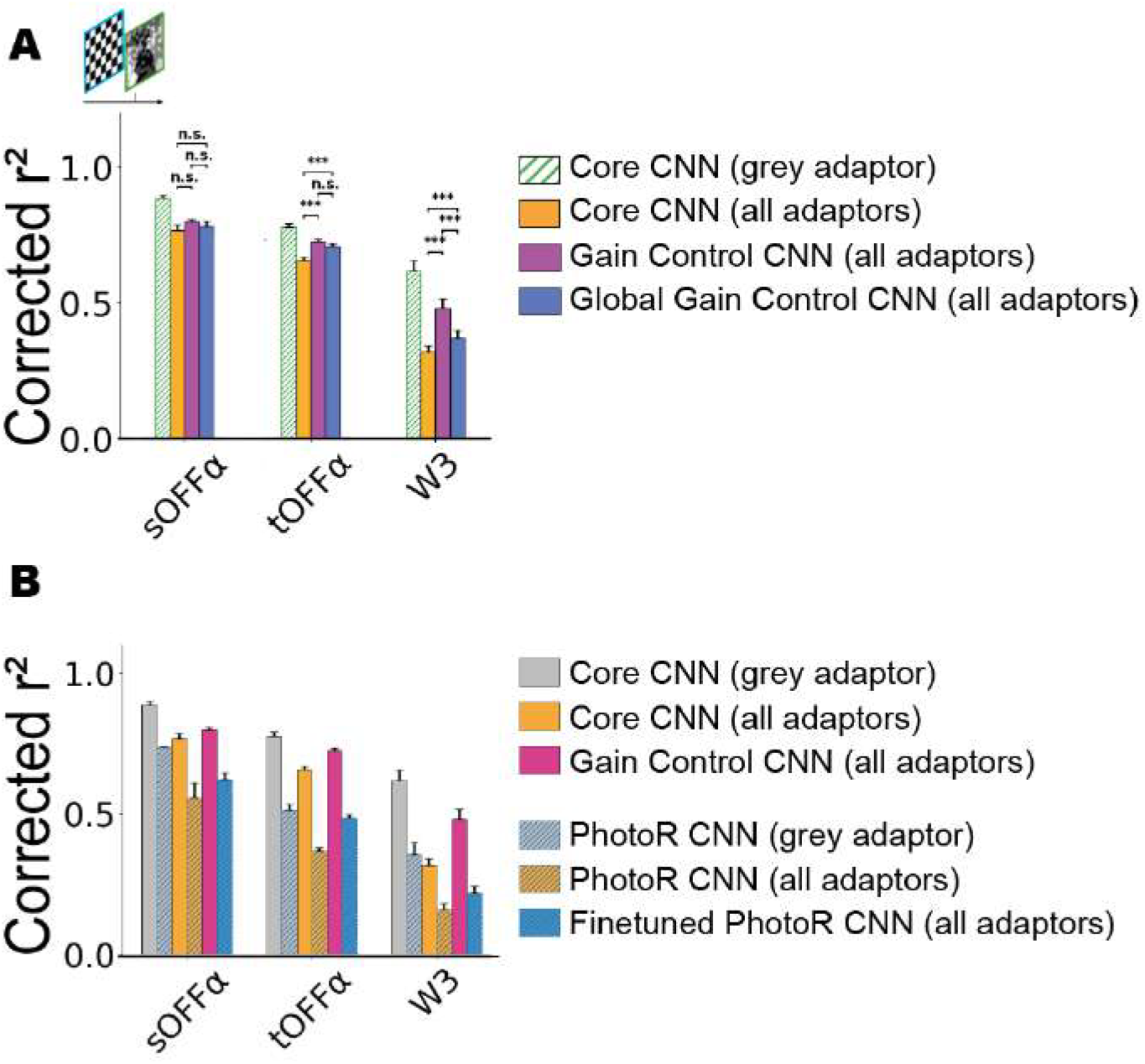
Alternative models of adaptation. **A**. Performance of the Core CNN, Gain Control CNN and Global Gain Control CNN (one gain control layer after the second non-linearity) at predicting responses to adaptor-probe sequences (grey and checkerboard adaptor, see Methods). Same cells as Fig. 2D. **B**. Same as A including performance of the Core CNN enhanced with a photoreceptor frontend layer (PhotoR model). All parameters of the PhotoR CNN were learnt using a grey adaptor only (In Distribution dataset). For the Finetuned PhotoR CNN, the biophysical photoreceptor layer parameters were finetuned using checkerboard adaptors (Out Of Distribution dataset).

**Supp. Fig.3.**
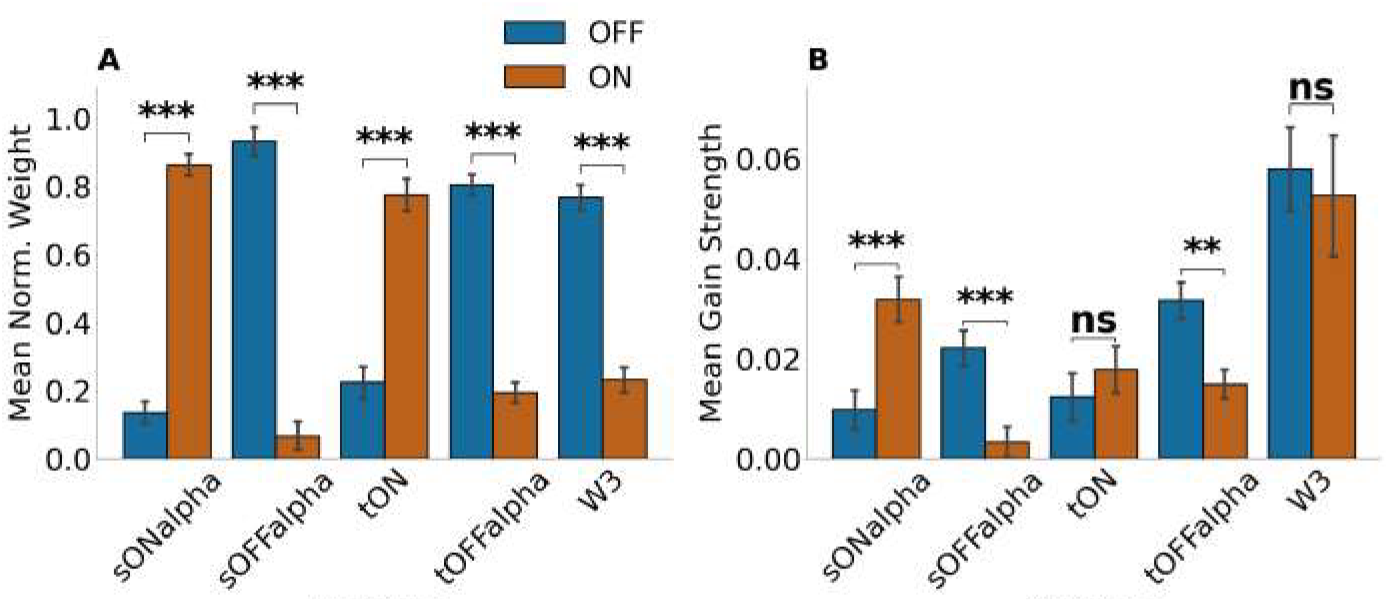
Distribution of subunit weights and gain control strength across retinal ganglion cell types. **A**. Mean normalized subunit weights for ON (red) and OFF (blue) polarities across five cell types: sONalpha (n = 26), sOFFalpha (n = 19), tON (n = 32), tOFFalpha (n = 15), and W3 (n = 43). Weights are normalized per cell to sum to 1 across all subunits. Bars represent the population mean, and error bars indicate the standard error of the mean (SEM). Statistical significance between ON and OFF weights was determined using a Wilcoxon paired signed-rank test. Weight distribution was significantly biased for all types (p < 1×10^−4^ for sONalpha, sOFFalpha, tON, tOFFalpha, and W3). **B**. Mean gain control strength (G, see Methods) for ON and OFF polarities across the same cell populations. Gain strength is derived from the temporal gain control layer of the model. Formatting, error bars, and statistical tests are identical to panel A. Gain strength differed significantly between polarities for sONalpha (p < 1×10^−4^), sOFFalpha (p = 0.0001), and tOFFalpha (p = 0.0037). No significant difference in gain strength was observed between ON and OFF subunits for tON (p = 0.0710) or W3 (p = 0.2134). All data are presented as mean and SEM.

